# Extensive nuclear gyration and pervasive non-genic transcription during primordial germ cell development in zebrafish

**DOI:** 10.1101/2020.01.10.901306

**Authors:** Stefan Redl, Antonio M. de Jesus Domingues, Stefanie Möckel, Willi Salvenmoser, Maria Mendez-Lago, René F. Ketting

## Abstract

Primordial germ cells (PGCs) are the precursors of germ cells, which migrate to the genital ridge during early development. Relatively little is known about PGCs after their migration. We studied this post-migratory stage using microscopy and sequencing techniques, and found that many PGC-specific genes, including genes known to induce PGC fate in the mouse, are only activated several days after migration. At this same timepoint, PGC nuclei become extremely gyrated, displaying general opening of chromatin and high levels of transcription. This is accompanied by changes in nuage morphology, expression of large loci, named PERLs, enriched for retro-transposons and piRNAs, and a rise in piRNA biogenesis signatures. Interestingly, no nuclear Piwi protein could be detected at any timepoint, indicating that the zebrafish piRNA pathway is fully cytoplasmic. Our data show that the post-migratory stage of zebrafish PGCs holds many cues to both germ cell fate establishment and piRNA pathway activation.

## INTRODUCTION

The specification of the primordial germ cells (PGCs) in the zebrafish starts at the 32 cell stage by the uptake of germ plasm, or nuage, an electron-dense, phase separated structure containing RNA and protein, into four blastomeres (Raz, 2003). The germ plasm-containing blastomeres initially divide asymmetrically and only daughter cells that inherit the germ plasm can at later stages become a germ cell (Knaut et al., 2000). At sphere stage the germ plasm disperses into smaller granules and associates increasingly with the nuclear membrane. From that point on, both daughter cells inherit germ plasm during cell divisions, and thereby the potential for becoming germ cells.

While the embryo develops, the PGCs, as defined by the presence of germ plasm, divide and migrate towards the genital ridge, where they arrive around 16-24 hours post fertilization (hpf) (Raz, 2003; Weidinger et al., 1999; Yoon et al., 1997). By this time, the number of PGCs has increased to 25-50 (Braat et al., 1999; Knaut et al., 2000; Weidinger et al., 1999; Yoon et al., 1997). Even though the initial specification and migration of PGCs in zebrafish is well studied, little is known about what happens to these cells from after their arrival at the genital ridge, until the time they start to form a definite gonad, which happens around 10 days post fertilization (dpf) (Leerberg et al., 2017; Tzung et al., 2014). The number of PGCs stays constant in this period, indicating these cells are not proliferating (Tzung et al., 2014). Furthermore, it was recently shown that global DNA methylation is maintained during this period, indicating that no de-methylated state occurs during the zebrafish germ cell development cycle (Ortega-Recalde et al., 2019).

The piRNA pathway is an RNAi-related pathway, found in the germ cells of many animals. Argonaute proteins are characteristic of all RNAi-related pathways, and represent the functional core of such pathways, by binding the small RNA co-factors and triggering gene-regulatory effects. PiRNAs are the small RNA-cofactors of a distinct branch of argonaute proteins, known as Piwi proteins. The zebrafish genome encodes two Piwi proteins: Ziwi and Zili (Houwing et al., 2007, 2008). Ziwi mainly binds piRNAs that are anti-sense with respect to their targets’ mRNA, which are mostly transposable elements (TEs), and are characterized by a strong bias for a 5’ uracil. In contrast, Zili binds mostly sense-piRNAs, which are enriched for an adenosine at position 10. Biogenesis of Zili-bound piRNAs is most likely triggered by Ziwi-mediated cleavage of a target RNA and vice versa. These relationships lead to a characteristic 10-nucleotide overlap, also named a ping-pong signature, between Ziwi- and Zili-bound piRNAs. In addition to this type of Piwi-directed piRNA biogenesis, another endonuclease, Zucchini (Zuc), also named Pld6, has been implicated in generating *Drosophila* piRNAs (Han et al., 2015; Mohn et al., 2015). In analogy to *Drosophila*, zebrafish Pld6 most likely generates Ziwi-bound piRNAs, and Zuc activity may be triggered by Zili-mediated cleavage of a transcript. Pld6 may generate several piRNAs from one RNA precursor through consecutive cleavage, resulting in so-called phased piRNAs, something that has been described for zebrafish as well (Gainetdinov et al., 2018).

Ziwi and its associated piRNAs are maternally provided via germ plasm, and are thought to provide a first line of defense against active transposon mRNAs in the zygote (Houwing et al., 2007). Maternal priming of the Piwi pathway has been observed also in *Drosophila* (Brennecke et al., 2008) and *C. elegans* (Luteijn et al., 2012; Shirayama et al., 2012), and in ciliates parental small RNAs guide the genome rearrangements of newly established zygotes (Lepere et al., 2008). These studies indicate that maternal priming of a small RNA pathway is a widely conserved phenomenon. Interestingly, maternally provided Ziwi protein can be preserved in germ cells for up to 3 weeks post fertilization (Houwing et al., 2007), and it was postulated that this pool may prime the zygotic piRNA pathway. *Ziwi* mutant germ cells are lost, starting at 20dpf (Houwing et al., 2007), suggesting that the phenotype may be triggered by loss of maternally provided Ziwi protein. The second PIWI protein, Zili, is not transmitted maternally, and only starts to be expressed zygotically around 3dpf (Houwing et al., 2008). While it first seems to be evenly distributed throughout the PGCs, and is only excluded from DAPI-dense regions, its distribution becomes increasingly restricted to perinuclear granules (Houwing et al., 2008). These observations have suggested that Zili may initially drive a nuclear piRNA pathway in the zebrafish, followed by restriction to the cytoplasm at later stages (Houwing et al., 2008). *Zili* mutant PGCs appear to arrest, but not to die, at a stage resembling PGCs at 3dpf, the time when Zili should start to be expressed (Houwing et al., 2007, 2008). This difference between *zili* and *ziwi* mutant PGCs is consistent with the idea that maternally provided Ziwi requires zygotically expressed Zili to sustain PGC development.

The piRNA biogenesis steps happen within an electron dense structure called nuage in germ cells. These structures represent phase separated condensates driven by both RNA and protein molecules. Nuage is found in germ cells of many animals, including mammals. Interestingly, different types of nuage have been described, based on different appearance in electron microscopy or the absence or presence of specific proteins (Aravin et al., 2009; Wan et al., 2018). Nuage in zebrafish adult germ cells is known to be very electron dense and tightly associated with mitochondria, while nuage in early embryos is more diffuse and less associated with mitochondria (Herpin et al., 2007; Huang et al., 2011). Proteins known to be required for normal nuage formation in zebrafish are Tdrd1 (Huang et al., 2011) and Tdrd6a (Roovers et al., 2018).

In an effort to better interpret mutant phenotypes in the zebrafish piRNA pathway, we set out to study the development of PGCs between 1 and 10dpf, by sequencing of small RNAs, mRNAs and total RNA populations, immunofluorescence and electron microscopy. The results not only detected zygotic activation of the piRNA pathway in this time-window, but also revealed that germ cell fate establishment may not take place until several days following arrival of the PGCs at the gonadal ridge. In addition, our work provides useful resources for future studies on zebrafish germ cell development.

## RESULTS

### Nuclei acquire a gyrated shape at 3dpf

The localization of the piRNA-pathway associated proteins Ziwi, Zili and Tdrd1 were already reported for 3dpf and 7dpf by our group. At the time we concluded that Ziwi and Tdrd1 localize to perinuclear granules at these timepoints, while Zili is both in the cytoplasm as well as within the nucleus at 3-7dpf, but becomes restricted to perinuclear granules at 7dpf. It was already noted that Zili was excluded from DAPI bright spots (Houwing et al., 2007, 2008; Huang et al., 2011). When we repeated these stainings, we noticed that it is actually extremely difficult to discern nucleus from cytoplasm at these stages. We therefore decided to use a LaminB1 antibody to stain for the nuclear envelope and combined it with stainings for Ziwi (Figure 1A). At 1dpf, PGCs had fairly round nuclei with large patches of Ziwi localizing perinuclearly (Figure 1A). However, at 3 and 6dpf we found the nuclear envelope to be heavily gyrated, forming very thin extensions at certain sites (Figure 1A, B). In some cells at 3dpf, Ziwi had a very punctate appearance (Figure 1C), but in others it was mostly diffuse throughout the cytoplasm (Figure 1A, B). Regardless of this, Ziwi always localized clearly outside the nucleus (Figure 1A,B).

**Figure 1:**
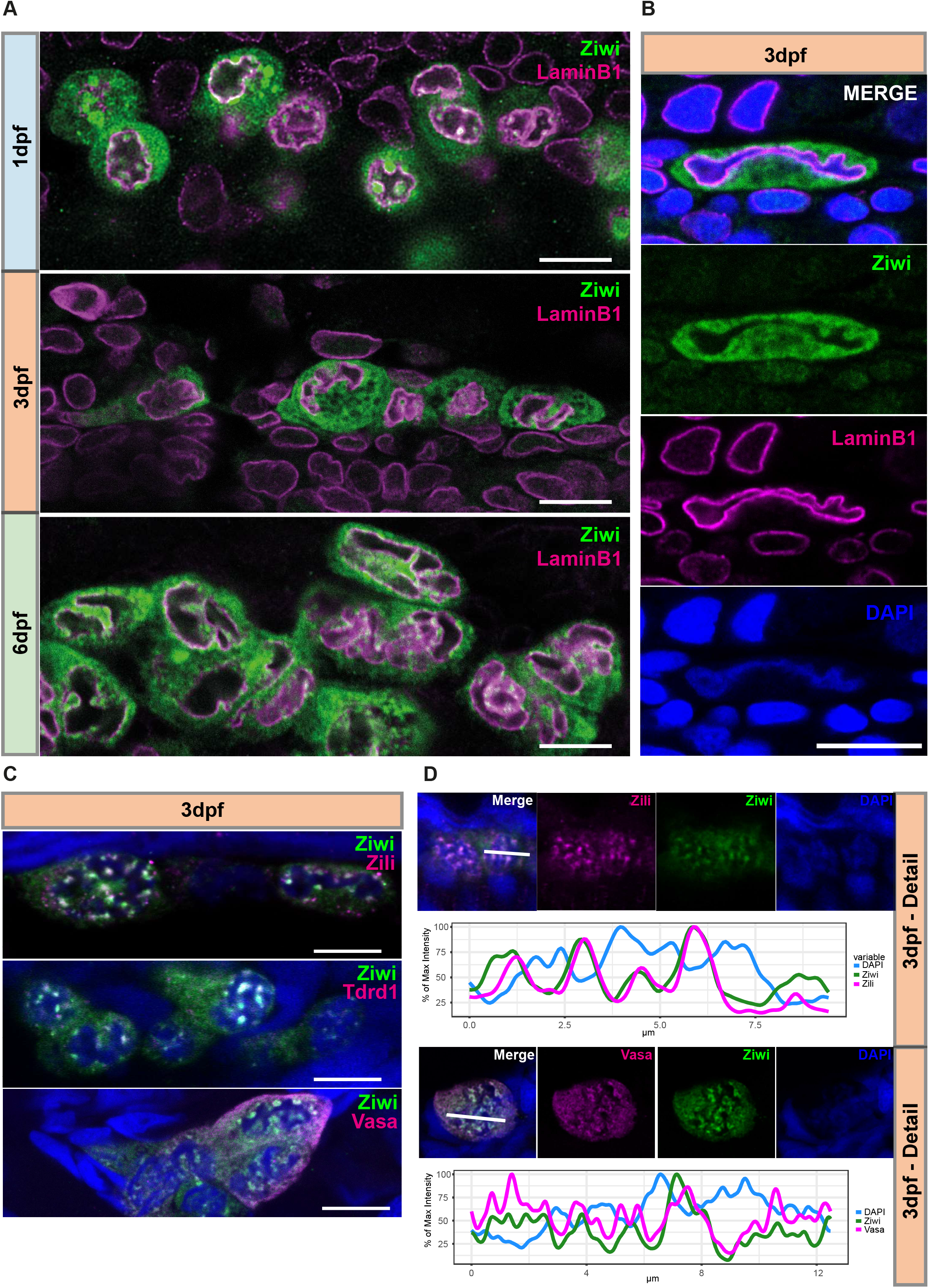
PiRNA pathway components localize outside of gyrated nuclei. **A:** Immune stainings for Ziwi (green) and LaminB1 (magenta) in PGCs at indicated timepoints. Scale bar 10μm **B:** Detail from a 3dpf PGC with Ziwi (green), LaminB1 (magenta) and DAPI (blue). Scale bar 10μm. **C:** Double-immune stainings for co-localisation of Ziwi and Zili, Tdrd1 and Vasa in PGCs at 3dpf. Scale bar 10μm **D:** Co-localisation analysis of Zili and Ziwi and Ziwi and Vasa. Blue: DAPI. Line plot of indicated selection for Ziwi and Zili (top) and Ziwi and Vasa (bottom) with DAPI. X-Axis: distance in μm. Y-Axis percentage of maximum intensity.

The peculiar nuclear morphology raised the possibility that the previously described nuclear localization of Zili (Houwing et al., 2008) in fact represented cytoplasmic pockets that are found seemingly inside the nuclei in optical sections. Unfortunately, the antibody-sources prevented direct LaminB1-Zili co-stainings to assess Zili localization. Hence, we combined Zili staining with DAPI and Ziwi to visualize both cytoplasm and chromatin (Figures 1C and D) at 3dpf, when Zili starts to be expressed. This revealed good overlap between Ziwi and Zili (Figure 1D), strongly suggesting that Zili is also cytoplasmic. Very similar outcomes were found for two additional piRNA pathway components, both known to be exclusively cytoplasmic: Vasa and Tdrd1 (Figures 1C and D). We conclude that both Ziwi and Zili localize to the cytoplasm at 3dpf, and that starting at 3dpf the PGC nuclei adopt heavily gyrated shapes, resulting in large cytoplasmic intrusions into the nuclei.

### Electron microscopy reveals different nuage types between developmental timepoints

To examine the observed nuclear morphology and other PGC characteristics in more detail, we turned to electron microscopy (EM). PGCs at 1dpf could be found in close proximity to the yolk syncytial layer (ysl) as reported previously (Braat et al., 1999). Interestingly, they did not contact the ysl directly but were separated from it by cellular extensions of somatic cells (Figure S1A). At all timepoints investigated we see that PGCs are closely associated with and surrounded by somatic cells (Figure S1A,B,C). The nuclei of 1dpf PGCs were mostly round with indentations found at the location of cytoplasmic nuage and sometimes showed a horseshoe shape, a characteristic also described for mouse PGCs (Clark and Eddy, 1975). Additionally, in agreement with previous reports (Huang et al., 2011) nuage was associated with clusters of nuclear pores (Figure 2A’). In contrast to nuage found in adult germline cells, in 1dpf PGCs the electron density of nuage had a rather granular appearance (Figures 2A and 2A‘). Hence, we will from here on refer to this type of nuage as ‘granular nuage’. Nuclei in 1dpf PGCs appear relatively euchromatic, but dark staining areas, indicative of heterochromatin could be easily detected.

**Figure 2:**
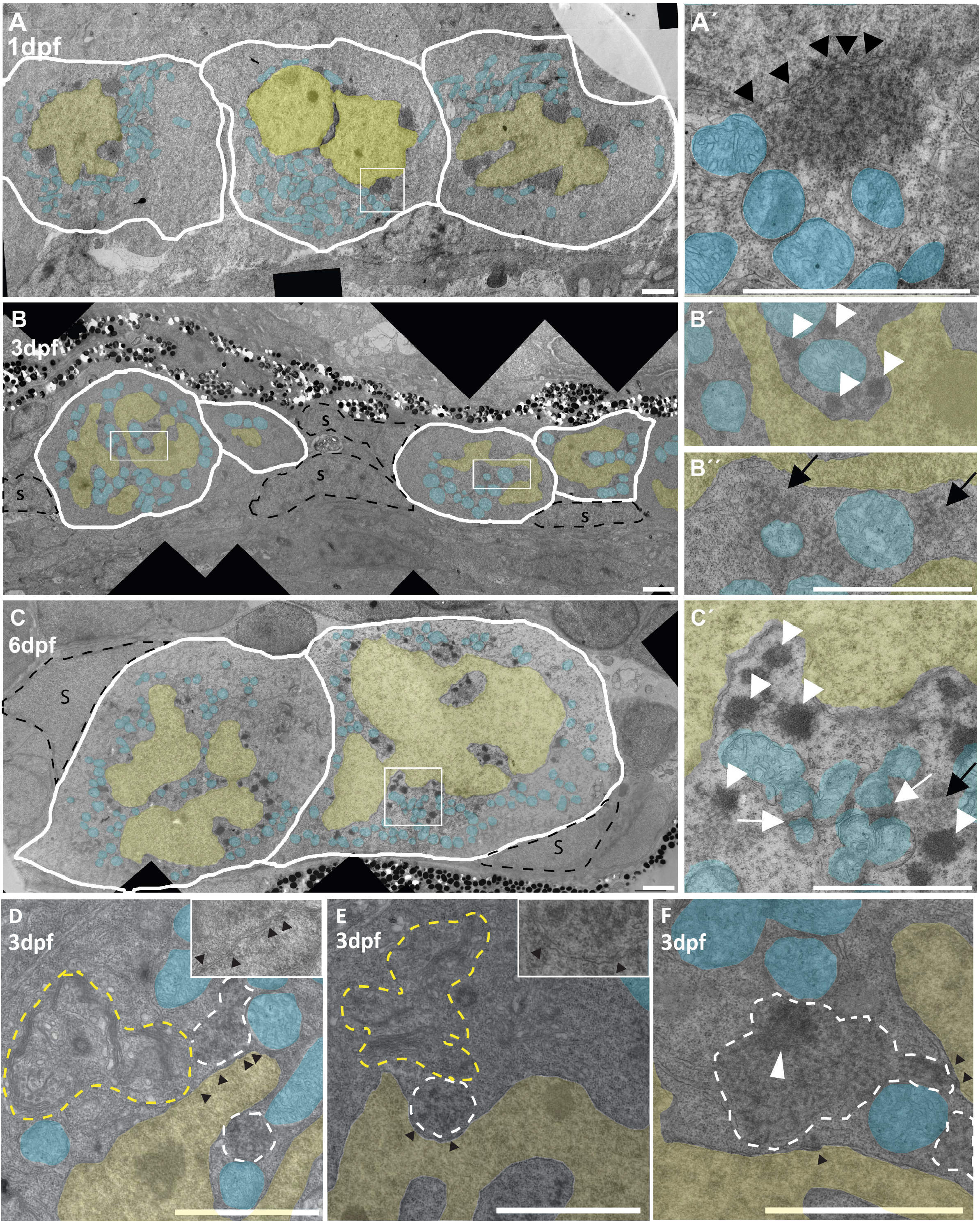
Electron microscopy on PGCs at different timepoints. **A:** 1dpf PGCs, nucleus (yellow) with sometimes one prominent invagination. Nuage can be found as perinuclear dark patches structures. **A’:** A zoom in on one nuage patch, showing a granular texture. Arrowheads point to nuclear pores. **B:** PGCs at 3dpf. The nucleus (yellow) has acquired an extremely irregular outline and nuage can be found with a granular texture (black arrow, detail in **B”**) or as dense granules (white arrowheads, detail in B”) around the nucleus. **C:** At 6dpf PGCs have grown in size and nuclei are still heavily gyrated. Nuage can be found perinuclearly (white arrowheads, black arrows) and between clusters of mitochondria (white arrows). Detail of **C** in **C‘**. Blue overlay: mitochondria. Yellow overlay: nucleus. S: marks somatic cells that contact PGCs extensively. **D** and **E**: Two examples of 3dpf PGCs, where fibrillar nuage (white dashed outline) is contacting nuclear pores (black arrowheads) and organelles, in this case mitochondria (blue overlay) and Golgi (yellow outline). Nuclei are marked with a yellow overlay. Insets show details without overlays. **F:** Example of granular nuage with a more compacted part; possibly a transition between granular and dense nuage. Scale bars: 2μm.

PGCs at 3 dpf were located adjacent to the pro-nephros (allentois) and a layer of pigment cells. Consistent with what we observed in immune fluorescence, PGCs had an extremely gyrated nucleus at 3dpf (Figure 2B). Nuclear invaginations were abundant, and cytoplasmic islands could be observed within nuclei in virtually any section. The nuclear membrane, however, was always intact. In addition, two types of nuage could be found. Some cells contained the granular nuage that we also observed in 1dpf PGCs (Figure 2B”). It appeared spread out, and was not only associated with nuclear pores but often also contacted organelles such as mitochondria and golgi (Figure 2D and E). The second type of nuage in 3dpf PGCs was more electron dense and more compact than the granular nuage (Figure 2B‘), resembling nuage as it is typically found in adult germ cells. We mostly saw one type of nuage per cell, suggesting these two types of nuage may reflect different developmental stages. In a few instances we could see nuage with both a granular and a dense area, possibly representing nuage in the process of switching (Figure 2F).

At 6dpf, PGC nuclei were still gyrated. The nucleoplasm contained very few heterochromatic (dark) patches and appeared more euchromatic compared to those observed in 3dpf PGCs. We quantified this using an approach that was recently published (Laue et al., 2019) and found that the level of heterochromatin at 6dpf was significantly lower than at 1 and 3dpf (Figures S1D and E). This suggests that chromatin globally becomes more open at this timepoint. Nuage could be found in numerous, uniformly dark staining patches all along the nuclear envelope, very similar in appearance to nuage found in adult germ cells (Figure 2C). Additionally, nuage in between mitochondria, detached from the nuclear membrane, was also abundant. Such structures have been named inter-mitochondrial cement in mammals (Eddy, 1974; Fawcett et al., 1970). It is, however, not clear if these are functionally distinct from nuage at the nuclear periphery. We note that some factors in the piRNA pathway, such as Zucchini, are associated with the mitochondrial membrane (Ipsaro et al., 2012; Nishimasu et al., 2012), and it seems likely that the mitochondria-nuage interactions are related to specific steps in the piRNA pathway involving these proteins.

The Tudor proteins Tdrd1 and Tdrd6a were shown to be necessary for either nuage or proper germ plasm formation in zebrafish: in *tdrd1* mutants nuage is almost completely disrupted in adult germ cells; while in *tdrd6a* mutants embryonic germ plasm is affected, but nuage seems unaffected (Huang et al., 2011; Roovers et al., 2018). Interestingly, Tdrd1 was found to only be expressed from 3dpf onwards, coinciding with the emergence of ‘adult-type’ nuage, while Tdrd6a was present in the maternally provided germ plasm that is found in the earlier stage PGCs (Huang et al., 2011; Roovers et al., 2018). We therefore probed the presence and localization of Tdrd6a in 1, 3, 6 and 10dpf-old PGCs (Figure S1F). Tdrd6a colocalized with Vasa and Ziwi in big perinuclear nuage granules at 1dpf, but in 3dpf PGCs was either colocalizing with Ziwi in perinuclear nuage or was absent from PGCs (* in Figure S1F, 3dpf). At 6dpf, Tdrd6a protein was no longer detected in PGCs, and only at 10 dpf a few germ cells showed re-appearance of Tdrd6a protein (Figure S1F). We hypothesize that cells with Tdrd6a correspond to cells displaying granular nuage in EM, and those lacking Tdrd6a to cells displaying adult-type nuage.

### PGCs show broad epigenetic changes between 1 and 10 dpf

We observed a change in appearance of chromatin during PGC development in electron micrographs (Figures S1D, E). This prompted us to investigate the histone modification status and RNA polII activity in the PGCs using immune stainings. First, we analyzed H3K9 trimethylation (H3K9me3), as a mark for constitutive heterochromatin. We found that, compared to surrounding somatic cells, H3K9me3 staining was relatively low in PGCs between 1 and 6dpf, and when present it was rather diffuse within the nuclei (Figure 3A). H3K9me3 only became detectable in the characteristic punctuate appearance in very few 10dpf PGCs (Figure 3A, arrowheads).

**Figure 3:**
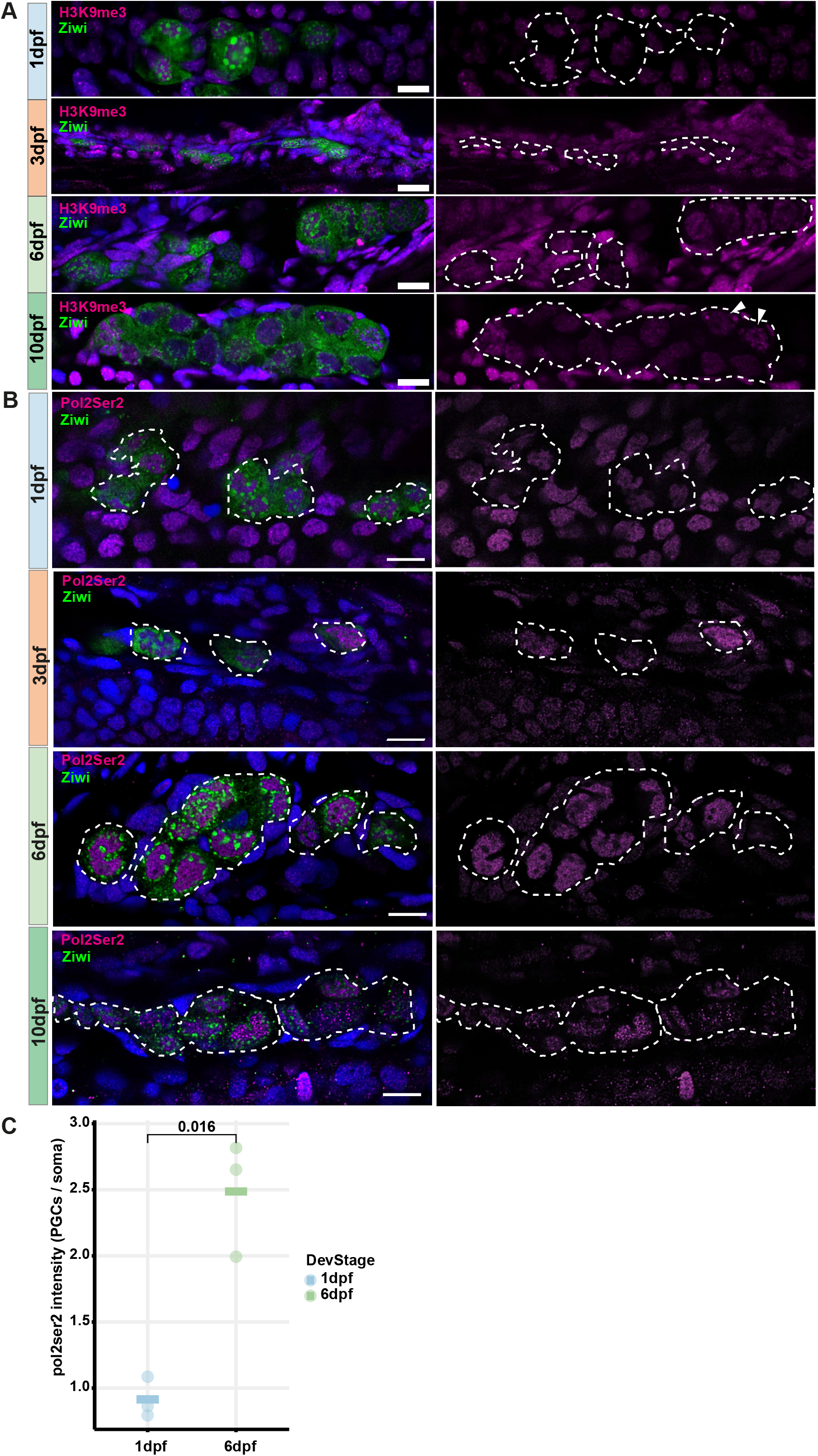
Histone modifications in PGCs. **A:** Double immunostainings for H3K9me3 (magenta) and Ziwi (green) at indicated timepoints. Scale bar: 10μm. **B:** Immune stainings for Ziwi (green) and Ser2-P modification in CTD of RNA polymerase II (Pol2Ser2) in PGCs at indicated timepoints. Right panel: single channels of Pol2Ser2 staining. Blue: DAPI. Scale bar: 10μm. **C:** Intensity ratios of Pol2Ser2 stainings between PGCs and somatic cells at 1 and 6dpf. Data points represent PGC/soma ratios from 3 images with a total of 30 PGCs per timepoint.

Next, we looked at H3K4me2 and H3K4me3, both associated with gene activity. Both marks were found to be present at all stages (Figure S2A, S2B), although H3K4me3 may be more abundant at 10dpf, specifically in nuclei with a round morphology (Figure S2B). Finally, we addressed polII activity levels by staining for elongating Pol2 (Ser2-P modification in CTD, Figure 3B). PGCs at 1dpf had levels of elongating Pol2 that are similar to the levels observed in their somatic neighbors. Starting in some cells at 3dpf, and in most, if not all cells at 6dpf, we observed very strong Pol2Ser2-P signals within the PGC nuclei, which appeared stronger than in the somatic neighboring cells. We analyzed staining intensities more closely in 1 and 6dpf PGCs by quantifying the intensity ratio between PGCs and surrounding somatic cells. This revealed a 2-3-fold increase in intensity between 1 and 6 dpf PGCs (Figure 3C). These data demonstrate that while PGCs are transcriptionally active already at 1dpf, this increases strongly towards 6dpf, where transcriptional activity appears to be very high in all PGCs. At 10dpf some cells still showed very high levels of Pol2Ser2-P, while others had reduced levels, with a more punctuate distribution, more consistent with somatic cells (Figure 3B).

### Germ line genes are upregulated from 3dpf onwards

To explore the transcriptional landscape of PGCs we created small RNA, total RNA and polyA selected RNA libraries, all originating from the same samples of FACS sorted PGCs (Figure S3A) and whole embryos at different timepoints in triplicate. We choose 0hpf (1 cell stage) to get an insight into what is maternally provided and then 1, 2, 3, 6 and 10dpf to analyze transcripts at different developmental stages. First, we will describe gene expression as obtained from the polyA-selected libraries.

To identify germline specific genes, we compared gene expression between ‘PGC’ and ‘total fish’ samples at matched time points. Using a stringent cut-off of false discovery rate (FDR) of 0.01 and a fold change increase of more than 30, we found 1600 genes that were specifically expressed in PGCs throughout the time course. Almost half of the genes were specific for one developmental timepoint (769/1600 genes), but we also found 131 PGC specific genes that were stably expressed in PGCs over the entire period. We called this set PGC-specific-stable-genes (PSGs) (Figure 4A and Table S1). The PSGs include all known piRNA pathway members and many known germline genes such as *zili*, *ziwi*, *tdrd1*, *tdrd9*, *vasa*, *tdrd6a*, *henmt1*, *rnf17*, *gtsf1*, *dnd1* and *dmrt1*. Additionally, a number of transcription factors like *zglp1*, *zgc:171506* and the TATA box binding protein like 2 (*tbpl2*) are among these genes.

**Figure 4:**
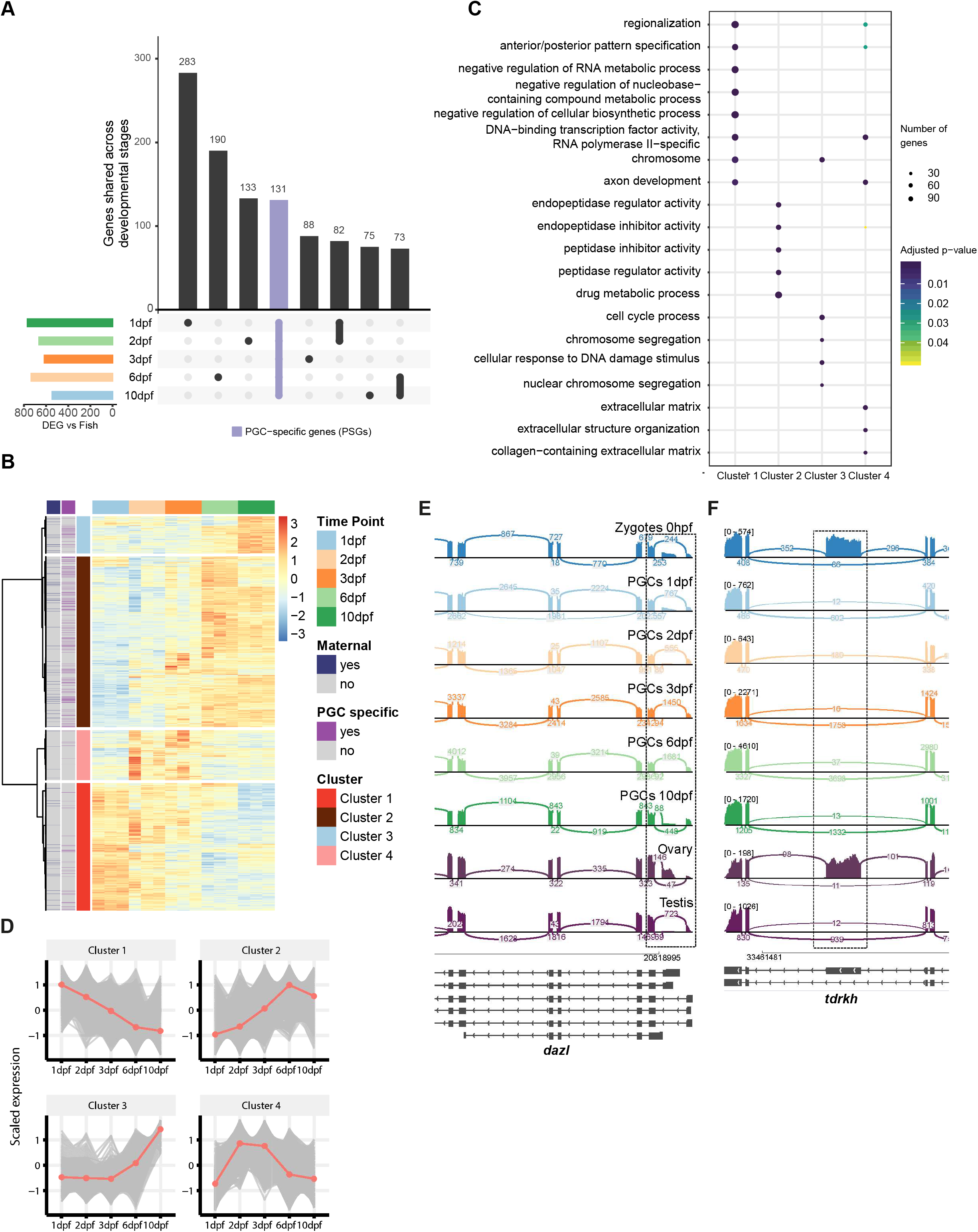
Gene expression analysis in PGCs. **A:** UpSet plot of PGC-specific genes. The colored bar plot on the left represents the number of germline enriched genes at indicated timepoints. 1, 2, 3, 6, 10dpf, as determined by comparing to ‘whole fish’ at the same timepoint. The vertical bars represent the number of PGC-specific genes present at certain timepoints (dots) or more than one timepoint (thick line). 131 genes are expressed at all 5 timepoint and enriched in the germline (purple): PGC-specific-stable-genes PSG. **B:** Hierarchical Clustering of genes differentially expressed in the PGCs. Maternally provided, annotation obtained from Aanes et al (2011), and PGC specific genes, as defined by our own data, are indicated in the left two columns. **C:** Gene ontology term enrichment of genes in the 4 different clusters. **D:** Average scaled expression (Z-score) of genes belonging to particular clusters. **E-F:** Sashimi plot depicting the alternative splicing of two germ-line genes, *dazl* and *tdrkh*. Dotted squares highlight the relevant splicing events. The values represent the total number of reads supporting the splicing events (sum of biological replicates).

We then investigated how genes that were expressed in PGCs, but that were not necessarily PGC specific, behaved throughout the time course. To do this we used hierarchical clustering to group genes into four clusters based on their expression at the different time points (Figure 4B and Table S2). In cluster 1 we found many genes whose transcripts are known to be maternally provided via the germ plasm, such as *granulito*, *nanos3*, *tdrd7a* and *ca15b* (Wang et al., 2013). As such, this cluster is characterized by genes whose expression declines during the developmental stages studied here.

Cluster 2 contains genes that are relatively lowly expressed at 1, 2, start to be noticeable at 3dpf and further increase in expression at 6 and 10dpf. This cluster contains many of the known germline genes and factors involved in the piRNA pathway. Previous studies showed that the expression of many piRNA pathway components start around 3dpf (Houwing et al., 2008; Huang et al., 2011). This is also true in our data-sets, but the effect at 3dpf is overshadowed by an even stronger increase in expression at 6dpf (Figure 4D).

Group 3 genes remain low in the first few days and then start to increase at around 6dpf, and continue to rise at 10dpf. These genes are associated with GO-terms including cell cycle process, (nuclear) chromosome segregation and cellular response to DNA damage stimulus (Figure 4C). These terms all fit well with the fact that at this time PGCs become mitotically active again (Leerberg et al., 2017). Other noteworthy genes in this cluster are *ddx6* and *dcp1a*. Both proteins are present in so-called piP-bodies in mouse. The piP-body in mouse does not only contain the Piwi protein MIWI2, the tudor domain protein TDRD9 and MAEL, but also classical components of P-bodies, namely GW182, XRN1 and the above mentioned DDX6 and DCP1a (Aravin et al., 2009). Possibly these observations are related to the changes in nuage we detected in our EM experiments (Figure 2). Cluster 3 also includes *kdm4aa*, a member of the JmjC family of histone demethylases, that in other organisms is involved in control of heterochromatin organization. It antagonizes H3K9 tri-methylation at pericentromeric heterochromatin in mammalian cells (Fodor et al., 2006; Klose et al., 2006). Expression of this gene could play a role in the observed loss of heterochromatin, described above.

Group 4 contains genes that are transiently expressed primarily at 2-3dpf. GO term analysis revealed that some of the genes are involved in RNA polymerase II transcription factor activity and sequence−specific DNA binding (Figure 4C). Interestingly *prdm1a* and *b*, two homologs of mouse *blimp1/prdm1*, a known transcriptional repressor required for germ cell development in mouse, also are part of this cluster. It is known that PGCs in zebrafish lose their germ cell fate if they do not reach the genital ridge (Gross-Thebing et al., 2017). It is possible that PGCs only become fully committed germ cells once they interact with the cells at the genital ridge at this point in development and that the group 4 genes play a role in this process.

### Splice variants expressed in PGCs

Whilst an in-depth investigation of alternative splicing and alternative use of transcriptional start sites in the context of PGC development is out of the scope of this work, we did notice interesting changes in some genes. For instance, *dazl* mRNA showed alternative usage of two distinct transcriptional start sites (TSSs) (Figure 4E, dotted box). While maternally provided transcripts were derived from both TSSs, the majority of transcripts produced in 2, 3, 6 and 10dpf PGCs stemmed from usage of only the most upstream TSS. Comparison with publicly available RNA datasets showed that the maternally provided transcript reflects the RNA expressed in ovary, while the transcripts produced in the PGCs resembled those expressed within the testis (Pasquier et al., 2016). *Tdrkh* is another example of a gene for which the maternal transcripts differed from those expressed in the PGCs (Figure 4F). Maternally provided *tdrkh* mRNA included a large exon in its 3’ part that is not present in PGC or testis mRNAs (Figure 4E, dotted box). This large exon contains four stretches marked as low complexity regions (Figure S3B and C). Such low complexity regions can play a role in phase separation (Hyman et al., 2014; Wheeler and Hyman, 2018), raising the option that these two different TdrKH variants may relate to the different types of nuage that we find during PGC development. Curiously, part of the low complexity region within the facultative exon may be sensitive to redox state (Figure S3D). Regarding the close association of TdrKH with mitochondria (Ding et al., 2019; Honda et al., 2013), this might mean that the TdrKH isoform containing the IDR could be influenced by the redox state of mitochondria.

### Transcription of large intergenic regions starting at 3dpf

Next, we analyzed rRNA-depleted libraries from the same timepoints for potential transcription producing non-poly-adenylated transcripts. Total RNA from the zygote was almost entirely deprived of intergenic regions, as expected by the lack of zygotic transcription (Figure 5A), and 1, 2 and 3dpf samples revealed slight increases in intergenic transcription in both PGCs and ‘whole fish’ samples. At 6dpf a striking increase in intergenic expression was observed in the PGC samples, and this further increases at 10dpf. This increase is not observed in ‘whole fish’ samples from the same timepoint, indicating that this is a PGC-specific effect. This is not due to a particular intergenic location since the number of intergenic loci in PGCs that show this trend also increases over time (Figure 5B).

**Figure 5:**
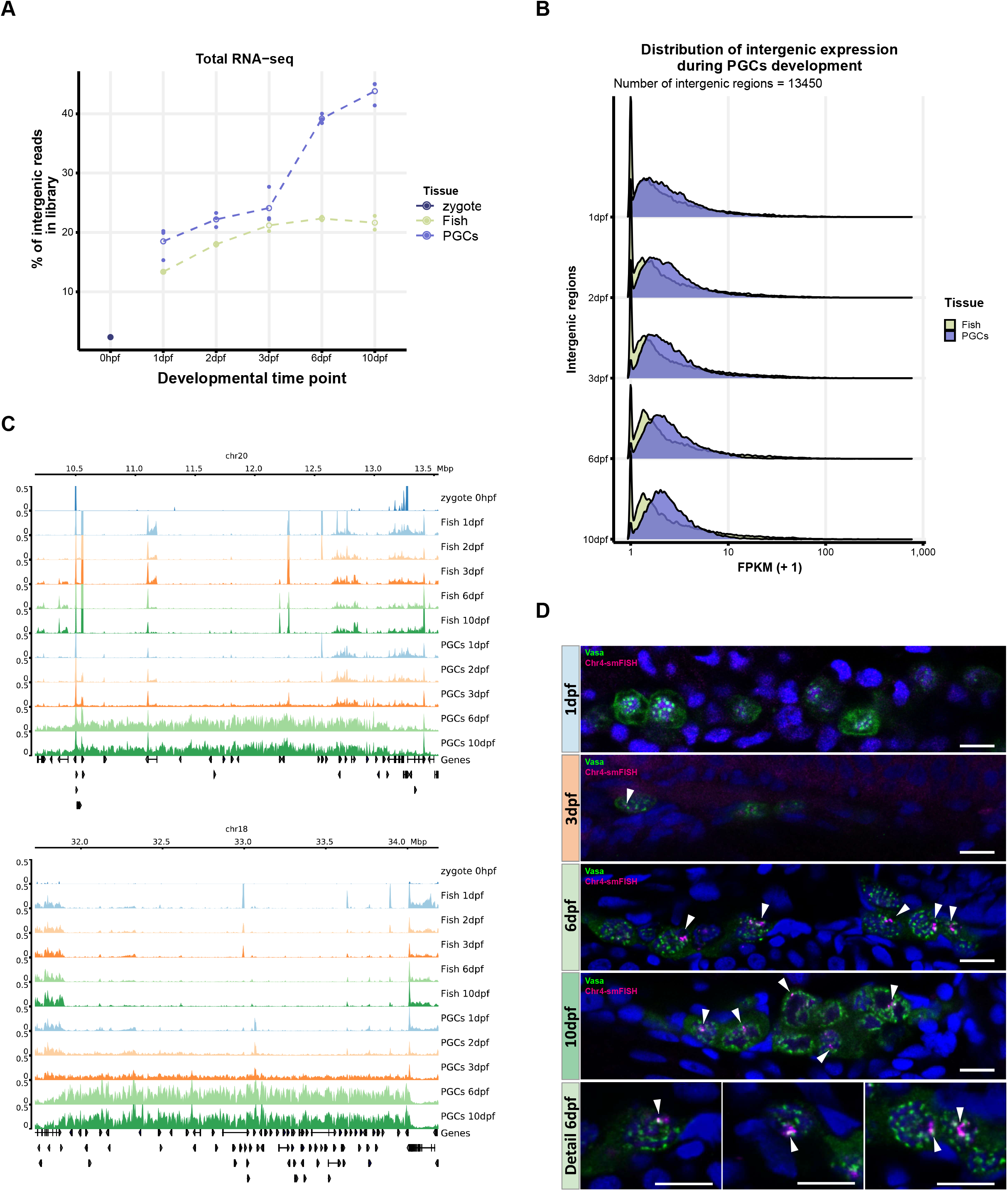
Expression of intergenic regions in the PGCs. **A:** Proportion of intergenic reads in total RNAseq libraries at the different indicated time points. **B:** Distribution of the expression of intergenic regions at the different timepoints. **C:** Example of two regions showing broad expression. **D:** smFISH using a probe designed to target transcripts derived from chr4: 48256970-48262902.

Visual inspection on a chromosomal scale suggested that starting at 3dpf, but more clearly visible at 6dpf and 10dpf, large intergenic regions start to get transcribed, specifically in the PGCs. We see this for instance for various regions including a 2 MB window on chromosome 18 and a 2,5 MB window on chromosome 20 (Figure 5C), as well as for the long arm of chromosome 4 (Figure S4A), which is relatively gene poor and is mostly transcriptionally silent (White et al., 2017), and is a major piRNA producing region in adult testis and ovary (Houwing et al., 2007). Interestingly, transcription from these regions comes from both strands, and while for a big part intergenic, also covers genes (Figure S4B-D). From here on we will refer to this type of expression as non-genic transcription, to reflect that it is neither excluded from nor limited to genes. To independently confirm expression, we designed smFISH probes against a region on chromosome 4 and used them to visualize expression of this locus throughout the time course. We did not see expression in either PGCs or somatic cells at 1dpf. At 3dpf we could detect smFISH signal in a restricted number of PGC nuclei and at 6 and 10dpf all PGCs were positive for the smFISH signal (arrowheads in Figure 5D). Like the RNAseq signal, the smFISH signal is restricted to PGCs. At a sub-cellular level, the signal was found within the nucleoplasm, overlapping with DAPI, but also in perinuclear granules that were positive for Vasa (Figure 5D). These findings are consistent with the possibility that these transcripts are exported to nuage, where they may be processed into piRNAs.

### PERLs are transposon-rich, piRNA producing loci

We next aimed to define loci that display dynamics as shown in Figure 5A in an unbiased manner. To do this, we identified large (>250kb) nearly contiguous regions of expression in both PGCs and ‘whole fish’ separately, and excluded those that are found in both these samples to isolate PGC-specific regions (for details see Methods). Finally, regions defined at different timepoints were combined to generate one set of loci, that we from now on will refer to as PERLs (PGC-Expressed non-coding RNA Loci). Visual inspection revealed that most of these PERLs indeed show the required characteristics, but we also noticed that certain PERLs may in fact be larger than called (example in figure S4A), and some might be missed due our fairly stringent definition. Nevertheless, we found that this set is as unbiased as possible and a fair representation of the broad expression phenomenon we noted in the PGCs. In total, we defined 201 PERLs (Table S3), deriving from all the zebrafish chromosomes (Figure 6A). We did not detect enrichment of PERLs on specific chromosomes, except for chromosome 4 (Figure 6A). Their size ranged between the implemented 250kb minimum and close to 3Mb (Figure 6B), and we detected a clear increase of PERL expression at 6dpf (Figure 6C).

**Figure 6:**
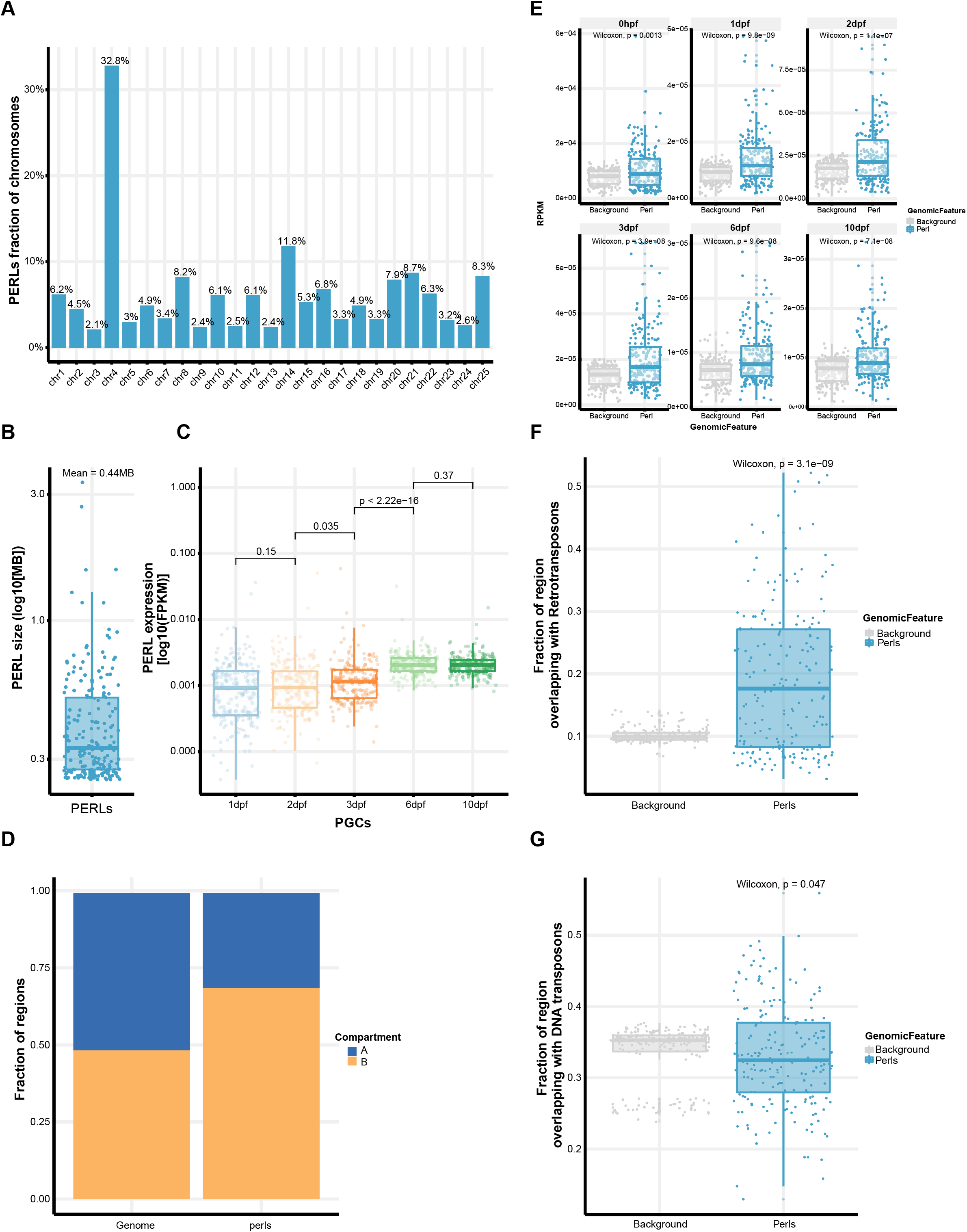
PERLs are transposon-rich, piRNA producing loci. **A:** PERLs as proportion of all zebrafish chromosomes. **B:** Size distribution of PERLs. **C:** Expression levels of PERLs across the different timepoints. Significance was tested with Wilcoxon test. **D:** Fraction of PERL base pairs that overlap with either A or B compartments. The number of genomic bases which belong to either of those compartments is also shown for comparison. **E-F:** Fraction of PERL bases which overlap with (E) RNA or (F) DNA transposons. Background is a set of size and chromosome matched random regions.

Given that heterochromatin visually disappears in the PGCs, and that heterochromatin has been proposed as a key driver in A/B compartmentalization (Falk et al., 2019) we checked whether PERLs may correlate with A or B status of chromatin as defined from HiC analysis on embryonic samples (Kaaij et al., 2018). Whilst the zebrafish genome is evenly split between A/B compartments, PERLs were over-represented in B compartments (Figure 6D).

Finally, we addressed the hypothesis that PERLs could be piRNA sources. To address this, we compared piRNA coverage (also see next section) in PERLs with that of a set of randomly selected genomic regions. Indeed, there was significantly higher piRNA expression from PERLs, and this phenomenon was consistently found in piRNA populations across development (Figure 6E). Interestingly, PERLs were also enriched significantly for retro-transposons, which are the major piRNA targets in zebrafish (Houwing et al., 2007; Kaaij et al., 2013), but not DNA transposons (Figure 6F and G). Given that DNA transposons cover a significantly larger fraction of the genome than retro-elements (Howe et al., 2013) this could indicate that retro-elements play a role in driving PERL expression.

### Evidence for zygotic piRNA pathway activation at 6dpf

Finally, we used the small RNA sequencing data to analyze piRNA dynamics between fertilization and 10dpf. Given the limited amounts of biological material, Ziwi and Zili IPs were not possible. In order to still enrich for bona-fide piRNAs over potential non-specific small RNA fragments, we concentrated on TE-derived small RNAs. These small RNAs displayed, at all time points, the characteristic Gaussian distribution with a peak at 27 nt (nucleotides) (Figure S5). From previously published work we know that the piRNA pool that is maternally provided via Ziwi is heavily anti-sense biased (Houwing et al., 2007), and this is what we also observed (Figure 7A). Interestingly, at 6 dpf this anti-sense bias significantly started to decrease, consistent with the idea that the piRNA amplification cycle is initiated and that Zili starts to be loaded with primarily sense piRNAs (Figure 7A). This was accompanied by an increase in ping-pong score (Figure 7B). These results are clear indications that the piRNA pool shifts from purely maternal piRNAs, to a situation where zygotic piRNAs are being produced.

**Figure 7:**
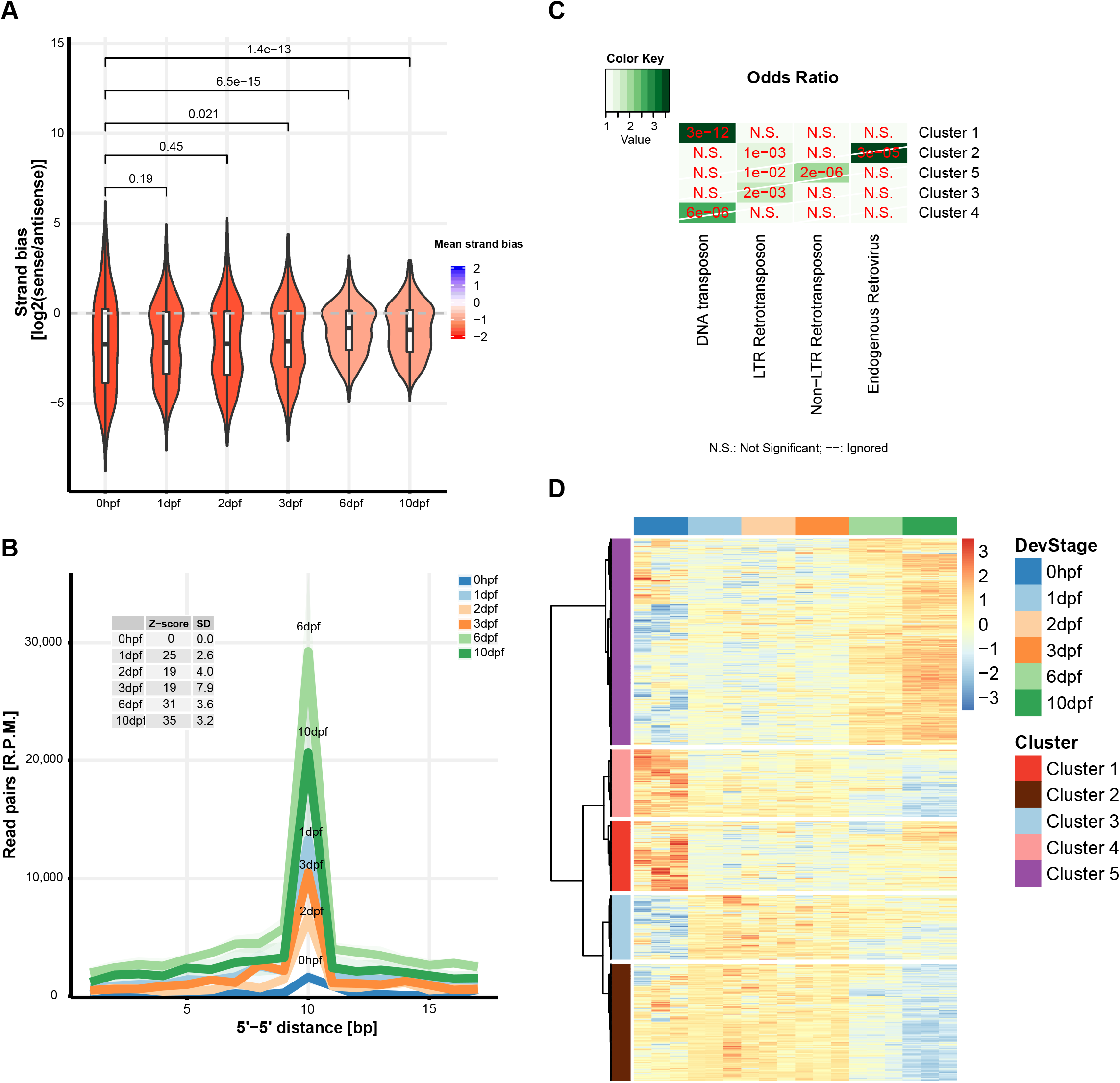
Zygotic piRNA pathway activity. **A:** Sense / Antisense bias of piRNAs increases during development (p-values generated using the Wilcoxon test). **B:** Ping-pong activity was assessed at the different time points by plotting 5’-5’distance of overlapping piRNAs from different strands. Z-scores were calculated to test significance. SD: standard deviation. **C:** Enrichment analysis of TE in the piRNAs target clusters in D. **D:** Hierarchical clustering of TE using the number of piRNAs mapping to those elements at the different time points.

We then investigated how piRNAs derived from different clades of TEs change throughout the time course. The piRNAs were grouped into 5 different clusters using hierarchical clustering and individual clusters were probed for enrichment of TE clades (Figure 7C). Cluster 5, which was enriched for LTR- and non-LTR-retrotransposons (p-value < 1e−02 2e−06 respectively, hypergeometric test, Figure 7C) strongly gained piRNAs at 6 and 10dpf (Figure 7D), consistent with the enrichment of retrotransposon enrichment within PERLs (Figure 6F). Indeed, these type of elements have been described before as major substrates in the zebrafish ping-pong cycle (Houwing et al., 2007, 2008).

## DISCUSSION

Following arrival at the genital ridge we found that zebrafish PGCs undergo many changes, both morphologically as well as molecularly. Our study reveals many interesting, thus far hidden characteristics in zebrafish germ cell development, providing many openings for further studies on zebrafish PGC development. Some of these aspects will be further discussed below.

### piRNA pathway activation in zebrafish PGCs

We described before that piRNAs, in particular those that are Ziwi-bound, are maternally provided to zebrafish embryos and can influence piRNA characteristics trans-generationally (Houwing et al., 2007; Kaaij et al., 2013). However, due to experimental difficulties in manipulating this maternal piRNA pool, the function of the maternal piRNAs in zebrafish is still unclear. Nevertheless, a number of potential functions, which are not mutually exclusive, can be thought of. For instance, maternally provided piRNAs may defend the early embryo against TE activity at stages before the zygotic piRNA pathway has been fully activated. The genome of zebrafish embryos undergoes several structural and epigenetic transitions during early development (White et al., 2017), and activation of TE activity can be expected to accompany such rewiring of gene expression. However, TE activity has not yet been specifically addressed at these stages of zebrafish development.

Studies in *Drosophila* have revealed that maternal piRNAs are very important in establishing a silencing response against TEs (Brennecke et al., 2008; Marie et al., 2016). Interestingly, this maternal effect in flies is thought to act not directly on active TEs, but through activation of piRNA clusters (Akkouche et al., 2017; Le Thomas et al., 2014). This happens through the nuclear piRNA branch that affects the chromatin structure of piRNA clusters in such a way that it stimulates piRNA production. Our work shows that the zebrafish most likely lacks such a nuclear piRNA mechanism, and hence, that maternal piRNAs are unlikely to directly affect the chromatin status of piRNA producing loci. One could therefore ask whether maternal piRNAs could still be directly involved in establishing the zygotic piRNA system in zebrafish? We hypothesize that maternal piRNAs in zebrafish indeed do that, simply by triggering the ping-pong amplification mechanism, as basically all required ping-pong-related proteins start to be expressed at the same point in time. The substrates for this process are likely the PERL-derived transcripts. Interestingly, *pld6* and *tdrKH* are activated at this same point in time as well, suggesting that the ping-pong mechanism may be tightly coupled to Pld6/Zucchini-driven piRNA production in zebrafish, as has been found in *Drosophila* (Han et al., 2015; Mohn et al., 2015). Indeed, signatures of this mechanism, that results in so-called phased piRNA production, have been found in zebrafish piRNA data (Gainetdinov et al., 2018). Finally, we note that absence of a nuclear piRNA branch is not zebrafish-specific. In many fish genomes only two Piwi proteins can be traced, suggesting that their piRNA systems may parallel that of zebrafish. Similarly, the cnidarian Hydra expresses only two Piwi proteins which both localize exclusively perinuclear and also in the silk moth *Bombyx mori* a nuclear piRNA pathway seems to be absent (Kawaoka et al., 2008; Lim et al., 2014).

### Nuclear gyration during PGC development

We describe exceptional nuclear folding of nuclei in PGCs. To our knowledge, such nuclear shapes have not been described before in healthy, normally developing, diploid cells. Somewhat similar nuclear shapes have been described in nuclei from cells of progeria patients, and in zebrafish mutants defective in a protease that processes LaminA (Tonoyama et al., 2018), but even in these cases the gyration was not as extensive as seen in the normally developing PGCs that we studied. How and why do the nuclei of the developing zebrafish PGCs become so gyrated? Since we often detect nuage structures within the cytoplasmic folds that extend into the nuclei, it is possible that deformations are triggered by strong RNA export forces acting on nuclear pores, or associated structures. The extensive intergenic transcription that we observed might provide such forces, as potentially very long piRNA precursors may at one end be still bound to the locus, or other nuclear structures, while at the other hand already become engaged by factors resident in nuage. Such binding of piRNA precursor transcripts has indeed been shown in *Drosophila* (Zhang et al., 2012).

The observed nuclear deformations in PGCs may also be linked to changes in genome ‘rigidity’, due to the loss of overall heterochromatin. Genomes are divided into compartments at different scales (Kim and Dekker, 2018), and phase-separation based mechanisms driven by such compartments have been implicated in nuclear organization (Gibson et al., 2019). Large scale loss of compartmentalization, due for instance to loss of heterochromatin (Falk et al., 2019), could lead to different mechanical properties of the genome and possibly more ‘fluid-like’ nuclear contents. Combined with strong transcriptional activity and export, this could trigger the observed nuclear shapes. Testing of such a hypothesis will require the identification of factors driving the loss of heterochromatin and whether such changes in chromatin state are indeed linked to changes in material properties of the nuclei.

We cannot tell whether the nuclear gyration bears functional relevance in itself. Possibly, the nuclear surface may need enlargement to accommodate high numbers of nuclear pores, which may in turn be needed for proper nuage development and piRNA functionality. But of course, the folding may only be a consequence, and have no functionality as such. Experiments targeting the functionality of nuclear gyration will likely need PGC-specific manipulation of gene expression between 3 and 10dpf. Currently such techniques are unfortunately not readily available for the zebrafish.

### Intergenic transcription during germ cell development in the mouse

Recently, a paper was published describing the opening of chromatin over large intergenic regions, named DADs, during mouse germ cell development, in particular in the male germ cell precursors, known as gonocytes (Yamanaka et al., 2019). Even though that paper did not address potential links between the opening of the chromatin, increased intergenic transcription and the piRNA pathway, it is certainly possible that what we see in zebrafish PGCs is mechanistically, and/or functionally related to the DADs in mouse gonocytes. Yamanaka et al. (2019) do not describe abnormalities in nuclear shape in gonocytes, at the time when their chromatin opens up.

### Establishment of germ cell fate in the zebrafish

How or when is germ cell fate established in zebrafish? While it has been well established that the maternally provided germ plasm is required to specify germ cells in zebrafish, it is yet unclear when the cells that inherit this germ plasm commit to germ cell development. Mis-localization of PGCs during their migration to the genital ridge can result in ectopic PGCs, and these cells have recently been shown to be able to take on somatic cell fates (Gross-Thebing et al., 2017). These results suggest that germ plasm inhibits somatic fates, rather than that it directly imposes germ cell fate. We describe activation of many ‘typical’ germ cell genes only at 3dpf and not before, and that at this time also homologs of genes known to drive germ cell fate establishment in mouse are transiently expressed. Examples of these genes are zebrafish homologs of mouse *prdm1* and *14* (*prdm1a* and *b*, and *prdm8*) and *ap2γ* (*tfap2a-e*), that are known to be required for expression of *nanos3*, *dazl* and *ddx4*/*vasa*. Interestingly, at this point in time PGCs are extensively enclosed by somatic cells. Possibly, these somatic cells represent the niches in which the PGCs receive the signals that trigger their further development into germ cells. The identification of molecular markers for these somatic cells will be required to help elucidate their role in germ cell development in zebrafish. We note that such relatively late establishment of germ cell fate may not be zebrafish specific, as also in mice, fixation of germ cell fate has recently been described to occur much later than was thus far assumed (Nicholls et al., 2019).

## ACKNOWLEDGEMENTS

We would like to thank the entire Ketting lab for fruitful discussions, especially Elke Roovers. Additionally, Yasmin El Sherif and Monika Kornovska are thanked for experimental support. We thank the Genomics, Flow Cytometry, Media Lab, Microscopy and Bioinformatics IMB Core Facilities for their contributions and valuable services.

## Methods

### Contact for Reagent and Resource Sharing

Further information and requests for resources and reagents should be directed to and will be fulfilled by the Lead Contact, René F. Ketting.

### Experimental Model and Subject Details

#### Zebrafish Lines

Zebrafish strains were housed at the Institute of Molecular Biology in Mainz and bred and maintained under standard conditions (26-28°C room and water temperature and lighting conditions in cycles of 14:10 hours light:dark) as described by (Westerfield, 1995). Larvae < 5 days post fertilization were kept in E3 medium (5 mM NaCl, 0.17 mM KCl, 0.33 mM CaCl_2_, 0.33 mM MgSO_4_) at 28°C. The *vasa:egfp* line was used for FACS sorting (Krøvel and Olsen, 2002). All experiments were conducted according to the European animal welfare law and approved and licensed by the ministry of Rhineland-Palatinate.

### Method Details

#### Immunostaining

Embryos were fixed in ice cold 4% PFA in PBS (pH 7.4) or in 80% Methanol/DMSO for 3h on Ice with gentle aggitation, washed with PBST (PBS and 0.1% Tween20), dehydrated in a standard methanol series and stored for at least 12 hours at −20°C. Followingly, wholemount staining was performed using a modified protocol from Inoue et al (Inoue and Wittbrodt, 2011).

Embryos were rehydrated from methanol and washed with PBSTw twice for 10 mins and 2 × 10min with PBS 1%Triton X-100. Followingly embryos older than 24hpf were decapitated and their tails were cut off for better penetrance of the solutions. Afterwards antigen retrieval was performed with 150mM Tris-HCl (pH 9) with 5 min RT incubation followed by 15 mins at 70°C with gentle agitation. After washings with PBSTw, embryos were put in blocking solution (PBSTw with 1% BSA, 10% sheep serum, 0.8% Triton-X100) for at least 90 mins followed by incubation with primary antibodies in blocking solution over night at 4°C. After extensive washing with PBSTw (6×30 mins), embryos were incubated with secondary antibodies o/n at 4°C. After another wash, DAPI was used to stain DNA. After additional washes, embryos weremounted ProLong™ Gold antifade mountant and imaged on a Leica SP5 confocal microscope with a 40X water objective (NA 1.3).

Immunostaining analysis were done as follows. Regions of interest (ROIs) were drawn for somatic cells and PGCs and the area, mean, min and max intensity levels of the Pol2Ser2-P channel were measured with ImageJ. Then the mean level of intensity between PGCs vs somatic cells was calculated and expressed as a ratio. Three measurements at each timepoints for somatic cells and PGCs were taken and ratios were compared between 1 and 6dpf. For the three datapoints, Pol2Ser2 intensities from 7 PGCs, 10 PGCs and 9 PGCs (1dpf) and 9 PGCs, 11 PGCs and 6 PGCs (6dpf) were compared to the intensities of somatic cells in the same image. Plotting and statistical testing was done in R using ggplot2 plugin (Wickham, 2016).

#### Confocal Imaging

Samples were imaged using a TCS SP5 Leica Confocal microscope using 40x Oil (NA of 1.3), 63x Oil (NA of 1.4) or a 40x Water (NA 1.2) objective. The following figures were deconvolved using Huygens software: Figure 1A, 1D, 3B, 5D.

#### Electron microscopy

Zebrafish embryos at indicated timepoints were killed on ice and immediately fixed with half-strength Karnovsky fixative (pH 7.4)(Karnovsky, 1965) and postfixed with 1% OsO4 in 0.1M Cacodylate buffer (pH7.4), dehydrated in an Acetone series and embedded in EPON. Semi-thin sections (1.5μm) were cut on a Reichert Ultracut 2040 and a Butler diamond knife (Diatome) until the desired area in the embryo was reached. Ultra-thin sections (90nm) were cut on a Reicher Ultracut E and collected on pioloform coated copper slot grids, dried and stained with leadcitrate under oxygen free conditions for 2 minutes. Sections were examined with a Zeiss LIBRA 120 and 2048×2048 pixel, 16 bit images were acquired with a Albert Tröndle Restlichtverstärker Systeme (TRS) ccd camera. Image analysis was performed using ImageJ. Nuclei were manually selected and electron-dense and non-electron-dense regions within nuclei were identified using thresholding. Following thresholding, particles with a minimum size of 0.03 μm2 were automatically selected and counted. Nucleoli were removed and for each image, the area of particles was expressed relative to the total nuclear area.

Values were collected and stored in Excel, and R ggplot2 plugin was used to create plots. Statistical testing was also done in R.

#### Single Molecule FISH

Single molecule FISH probes were designed against an intergenic region on chromosome 4 (danRer10 Chr4: 48257970-48262902) with Stellaris probe designer with 48 probes targetting a region spanning 5933 bases. Fixed, dehydrated and then rehydrated embryos (tail and head was cut off from embryos) were incubated in hybridisation buffer for 30 mins at 30°C. Primary antibody and Stellaris probes (125nM probe) were hybridised in hybridisation buffer o/n at 30°C. After incubation samples were washed with wash buffer at 30°C twice while secondary antibody and DAPI were added during the second wash step (45mins). After another wash step, embryos were mounted in anti-bleach medium (GLOX buffer containing Trolox, Catalase and Glucose oxidase see recipies and chemicals section)

#### Protein sequence analysis

Protein sequence analysis for facultative TdrKH exon was performed using IUPred2 with long disorder prediction and the use of the context-dependent prediction ANCHOR2 and the (experimental) redox-state plugin (Mészáros et al., 2018).

#### FACS sorting

The *vasa:eGFP* line (Krøvel and Olsen, 2002) was used to FACS sort PGCs at different timepoints. Embryos were collected and incubated with TrypLE™ Express for 40 to 70 minutes, killed on ice and gently pipeted up and down with a glass pipet and/or a 200μl low retention pipet tip. After visual inspection, cell suspension was separated from trunks using a 100 μm sieve. Following another 5-15 minutes of digestion, FCS was added to 10% of total volume. Cells were spun down at 500g for 5min at RT, washed with PBS, resuspended in PBS with 2% FCS, put on Ice and immediately subjected to FACS using a 85μm nozzle on a BD FACSAria III SORP (Becton Dickinson). 1000 to 2500 cells were sorted directly into Trizol. RNA from sorted PGCs and whole embryos was extracted with Trizol and stored in MQ at −80°C until library preparation was done.

#### NGS library preparation

##### Poly-A RNA seq

NGS library prep was performed with SmartSeq2 RNA-Seq System following NuGen`s standard protocol (M01406v2). Libraries were prepared with a starting amount of 1 ng and amplified in 12 PCR cycles.

Libraries were profiled in a High Sensitivity DNA on a 2100 Bioanalyzer (Agilent technologies) and quantified using the Qubit dsDNA HS Assay Kit, in a Qubit 2.0 Fluorometer (Life technologies).

Samples were pooled in equimolar ratio and sequenced PE for 2x 75 cycles plus 16 cycles for the index read.

##### Total RNA sequencing

NGS library prep was performed with NuGen Ovation SoLo RNA-Seq System following NuGen`s standard protocol (M01406v2). Libraries were prepared with a starting amount of 1 ng and amplified in 14 PCR cycles.

Libraries were profiled in a High Sensitivity DNA on a 2100 Bioanalyzer (Agilent technologies) and quantified using the Qubit dsDNA HS Assay Kit, in a Qubit 2.0 Fluorometer (Life technologies).

Samples were pooled in equimolar ratio and sequenced PE for 2x 75 cycles plus 16 cycles for the index read. Small RNA sequencing

##### Small RNA sequencing

NGS library prep was performed with NEXTflex Small RNA-Seq Kit V3 following Step A to Step G of Bioo Scientific`s standard protocol (V16.06). Libraries were prepared with a starting amount of 1ng and amplified in 25 PCR cycles.

Amplified libraries were purified by running an 8% TBE gel and size-selected for 18 – 40nt. Libraries were profiled in a High Sensitivity DNA on a 2100 Bioanalyzer (Agilent technologies) and quantified using the Qubit dsDNA HS Assay Kit, in a Qubit 2.0 Fluorometer (Life technologies).

Samples from 1, 3 and 6dpf and samples for zygote and 10dpf PGCs were pooled in equimolar ratio and sequenced on 2 NextSeq 500 Flowcell, PE for 2× 75 cycles plus 16 cycles for the index read.

#### Data processing and analysis

##### smallRNA-seq

Raw reads were checked for quality with FastQC before adapter trimming with cutadapt (v1.9.1) -a AGATCGGAAGAGCACACGTCT -O 8 -m 21 -M 51, followed by removal of sequences with low-quality calls using fastq_quality_filter (-q 20 -p 100 -Q 33) from the FASTX-Toolkit (v0.0.14). PCR duplicates were removed making use of UMIs added during library preparation by collapsing reads with the same sequence, including UMIs, using a combination of unix command-line programs. Processed reads were mapped to the Zv10 Danio rerio genome assembly with bowtie (v0.12.8) -q --sam --phred33-quals --tryhard --best --strata --chunkmbs 256 -v 2 -M 1.

To match each mapped read, that is, small RNA, with a genomic element, we downloaded repeat element annotations the UCSC genome browser track RepeatMasker (Zv10) and gene annotations from iGenomes (Danio rerio, GRCz10). We then intersected mapped reads with these annotations with bedtools intersect -wa -wb -bed -f 1.0 with the either the flags -s or -S to determine which small RNAs map sense or antisense, respectively, the annotated features. Length profiles were obtained by filtering reads mapping sense or antisense to genetic elements annotated as belonging to a particular transposon class / biotype, and summarizing their length.

Ping-pong signal was determined by calculating the base pair distance between the 5’ of piRNAs overlapping in opposite strands (Brennecke et al., 2007). The Z-score was calculated as in Wasik et al (Wasik et al., 2015), and is defined as Z = (P10-M)/SD, where P10 is the number of reads pairs with 10 bp overlap, and M and S are the mean and standard deviation, respectively, of the number of read pairs at with 1-9 and 11–30 overlap. Sense / Antisense bias was determined by calculating the ratio of reads mapping in the same or the opposite strand for each annotated transposable elements (DNA or RNA in repeatmasker). To generated scatterplots comparing piRNA abundances in two conditions, small RNAs that mapped to transposable elements were quantified per element and normalized to mapped reads.

Nucleotide bias of piRNAs determined by summarizing the number of time a base is present in any given piRNA position. For the downstream U bias, the downstream annotated genomic bases, in the same mapping strand of the piRNA, were used (Han et al., 2015; Mohn et al., 2015).

TE targeting was determined by estimating the number of piRNAs reads mapping to repeat elements using repEnrich v1.2 with default settings (Criscione et al., 2014). repEnrich uses a two-step approach mapping to the genome and to transposable element sequences, to utilize both uniquely and mulimappping and accurately infer read counts on repetitive elements.

Targeted TEs were clustered using hierarchical clustering in R [hclust(as.dist(1-cor(t(rldLRT_05), method=“pearson”)), method=“ward.D”)].

##### mRNA-seq and total RNA

mRNA and total RNA read processing and mapping library quality was assessed with FastQC before being aligned against the D. rerio genome assembly and the genome annotation (Zv10 / GRCz10) with STAR v2.5.2b (–runMode alignReads –outStd SAM –outSAMattributes Standard –outSAMunmapped Within –outSJfilterReads Unique – outFilterMismatchNoverLmax 0.04 –clip3pAdapterSeq CTGTCTCTTATACACATCT –sjdbGTFfile–sjdbOverhang 82). Read alignment was performed using the full genomes and reads mapping to unassembled contigs were removed after read alignment. mRNA Coverage tracks were generated with deepTools v2.4.3 (bamCoverage –smoothLength 60 –binSize 20 – normalizeUsingRPKM) (Ramírez et al., 2016), and total RNA coverage tracks were created with bamCoverage --binSize 50 --smoothLength 500 --normalizeUsing CPM --blackListFileName blacklisted.bed --effectiveGenomeSize 1369631918. blacklisted.bed is available as supplemental data. Stranded coverage tracks were created as above but deepTools’s code was modified to allow for normalized of stranded signal to total mapped reads.

Public datasets used to for ovary and testis sashimi plots was obtained from SRR1524248 and SRR1524249, and processed as described above.

##### Gene and transposon expression

Differential expression and clustering Gene / transcript expression was quantified with salmon v0.11.0 (Patro et al., 2017) (salmon quant -l A -1) or salmonTE v0.2 (Jeong et al., 2017) for the transposable elements expression (SalmonTE.py quant –reference=dr –num_threads=4 – exprtype=count). Differential expression analysis were performed with the Bioconductor package DESeq2 v.1.18.1 (love_moderated_2014). The following settings were used to determine differentially expressed genes, depending on the goal: (i) to determine which genes are changed during PGC development at any time point, we defined a reduced model in combination with a likelihood ratio test [DESeq(dds, test=“LRT”, reduced= ~ 1)]; whereas to determine which genes are PGC-enriched a simple pairwise comparison fish vs PGCs at matched time points was executed. Genes were clustered using hierarchical clustering in R [hclust(as.dist(1-cor(t(rldLRT_05), method=“pearson”)), method=“ward.D”)]. The list of maternally contributed genes was obtained from Aanes et al (2011), supplementary file 1A. To generate a list of genes that are very likely PGC-specific, only genes with fc >= 30 enriched in the PGCs (vs fish) and an fdr < 0.01 were selected. These cut-offs were based on the most stringent threshold that would still be able to retain well know PGC-specific genes. Normalized counts were obtained from DESeq2. Go term enrichment analysis for categories was performed with the Bioconductor package ClusterProfiler (Yu et al., 2012), using all genes with at least one count as the background set.

TE expression was quantified with salmonTE v0.11.0, SalmonTE.py quant --reference=dr --exprtype=count, that uses salmon to quantify TE and summarize according to TE element (Jeong et al., 2017). Hierarchical clustering was performed as described above for gene expression.

##### Intergenic expression

Intergenic regions were defined by extending any annotated region in the GRCz10 ensembl GTF by 2500 bp and then taking the complementary genomic locations (bedtools slop -i stdin -g chrom.sizes -b 2500 | bedtools merge | bedtools complement). Non-annotated rRNA locations identified with blast (Table S4) and other contaminants were also discarded. Reads mapping to intergenic regions were counted with bedtools coverage -split -counts -F 1.0.

##### PERL identification

We identified the PERLs using a stepwise approach guided by our observations of large contiguous genomic region being expressed at certain time points, which included both genic and non-genic regions: (i) we used definder to call expressed regions separately for each time point and tissue (average coverage of the replicates). Derfinder calculates the mean (normalized) coverage at every genomic base. Bases passing the cuttoff (0.25) and within 3000bp of each are then joined in a single region. (ii) ERs that are close to each other by less than 3 kb are merged in a single contiguous expression cluster. (iii) To identify only those regions specific to PGCs, at each time point the PGC clusters were overlapped with those of the Fish, and those that overlap by more than 10 % of (PGC cluster) bases were excluded. (iv) Clusters smaller than 250kb are removed to keep only large PGC-specific expressed regions or PERLs. Finally, since the observed global expression tends to take place at later developmental stages (3-6), we excluded those regions also present in at 1-2 dpf.

##### PERL characterization

Unless stated otherwise, as a background for comparisons we created 100 sets of background regions matched for size and chromosome with the PERL regions. These background regions were generated with bedtools shuffle -i perls.bed -g -excl perls.bed -chrom. To estimate PERL coverage/expression, reads mapping to each PERL were counted with bedtools coverage -split -counts -F 1.0, normalized to size and library depth (RPKM) and the expression for each replicate averaged. piRNA abundance in PERLs was determined using 23-33 small RNA reads with multiBigwigSummary BED-file. Coverage was then normalized for size and the biological replicates averaged, or in the case of the background regions the matched PERL regions were averaged. Statistical comparisons were done with the Wilcoxon rank test. The base overlap with transposons and A/B compartments was determined with coverageBed (fraction of covered bases).

##### Data deposition

All newly generated NGS data were deposited at NCBI, BioProjectID PRJNA597223.

