## Supplemental figures and legends for "Extensive nuclear gyration and pervasive non-genic transcription during primordial germ cell development in zebrafish"

Zebrafish; primordial germ cell; zygotic activation; piRNA; intergenic transcription; nuclear morphology; gyration; nuage

#### Supplemental figure legends

**Figure S1** Electron micrographs of PGCs

**A, B** and **C** show electron micrographs of PGCs at 1, 3 and 6 dpf respectively. Yellow overlay shows nucleus, green overlay marks somatic cells that are in close association with the PGCs, magenta overlay marks yolk syncytial layer (ysl). **D**: representative images of thresholding for heterochromatin analysis in PGCs. * marks a nucleolus, which was excluded from heterochromatin analysis and nuclear area. **E**: Dotplot showing heterochromatin in percentage of total nuclear area in PGCs at indicated timepoints. **F**: Double immune stainings in 1, 3, 6 and 10dpf PGCs for Ziwi (green) and Tdrd6a (magenta). Scale bars EM images: 2µm. Scale bars immune stainings: 10µm.

**Figure S2** Histone modifications in PGCs

**A:** Double immune stainings for Histone modification H3K4me2 (magenta) and Ziwi (green) in PGCs at indicated timepoints. **B:** H3K4me3 (magenta), Ziwi (green) double immune staining at indicated timepoints. Blue: DAPI. Scale Bar 10µm.

**Figure S3:** An alternative exon in *tdrkh* possesses 4 disordered regions

**A:** Representative FACS plot of *vasa-eGFP* positive PGCs. The boxed region reflects the sorted cells used for the analyses of PGCs. **B:** Amino acid sequence encoded by the *tdrkh* facultative exon. Predicted disordered regions (uniprot) are underlined. **C:** Disorder prediction of the facultative exon with IUPred2A (red line). Scores above the line reflect likelihood of being disordered regions. Regions correspond to sequences identified as disordered regions in uniprot. Blue line shows ANCHOR2 prediction. ANCHOR2 tries to identify protein sequences where protein binding partners can initiate the transition between a structured and unstructured state. **D:** Using context dependent predictions, a redox-sensitive region is identified that makes a larger region disordered, depending on being in a folded or unfolded state. These redox-sensitive regions can play a role in protein function in different subcellular conditions (Erdős et al., 2019; Reichmann et al., 2018).

**Figure S4:** Examples of PERLs

**A:** Coverage tracks showing the right arm of chromosome 4 and total RNA coverage at the different timepoints. Locations of PERLs in this region are also shown. **B:** Stranded coverage tracks of the same region of chromosome 18 shown in Figure 5C. **C:** Stranded coverage tracks of the same region of chromosome 20 shown in Figure 5C. **D:** Stranded coverage tracks of right arm of chromosome 4.

**Figure S5:** piRNAs show the typical length profile throughout the time course

Length distribution of small RNAs reads that map sense (blue line) or anti-sense (red line) to TEs per time point in PGCs. Lighter shading indicates the standard deviation of three biological replicates.

**Table S1:** PSG genes.

This tab-delimited file contains all the genes we marked as stably and specifically expressed in PGCs.

**Table S2:** IDs of genes linked to the four clusters we defined.

This tab-delimited file provides the information on which genes are in which cluster.

**Table S3:** PERLs.

This bed file contains the annotation for all the PERLs we identified.

**Table S4:** Non-annotated rRNA loci.

This bed file contains the annotations of non-annotated rRNA loci (see Methods).

Figure S1

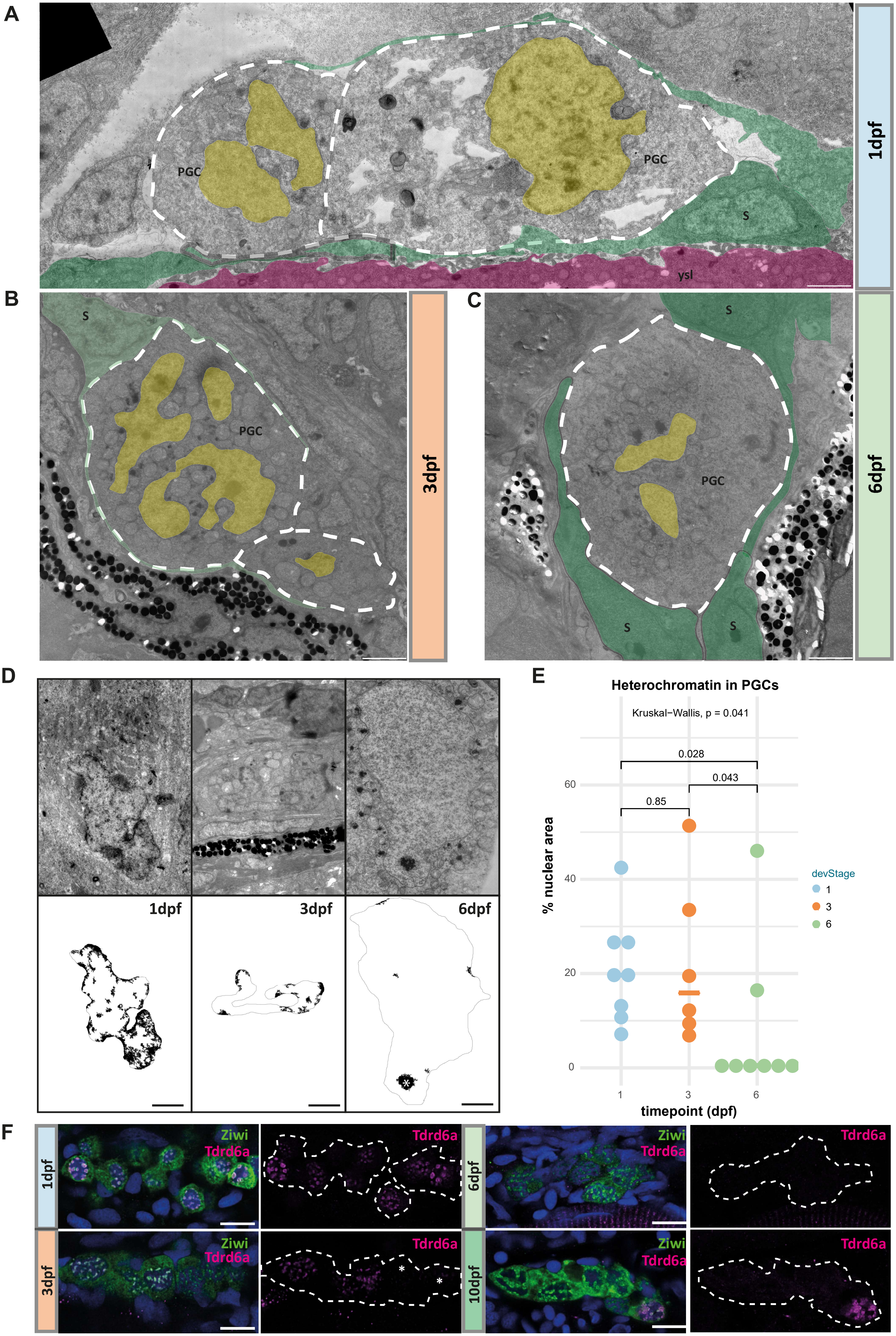
A
PGC
PGC
S
ysl
1dpf
B
S
PGC
3dpf
C
S
PGC
S
S
6dpf
D
1dpf
3dpf
6dpf
E
Heterochromatin in PGCs
Kruskal−Wallis, p = 0.041
0.028
0.043
0.85
% nuclear area
60
40
20
0
1
3
6
timepoint (dpf)
devStage
1
3
6
F
1dpf
Ziwi
Tdrd6a
Tdrd6a
6dpf
Ziwi
Tdrd6a
Tdrd6a
3dpf
Ziwi
Tdrd6a
Tdrd6a
10dpf
Ziwi
Tdrd6a
Tdrd6a

### Figure S2

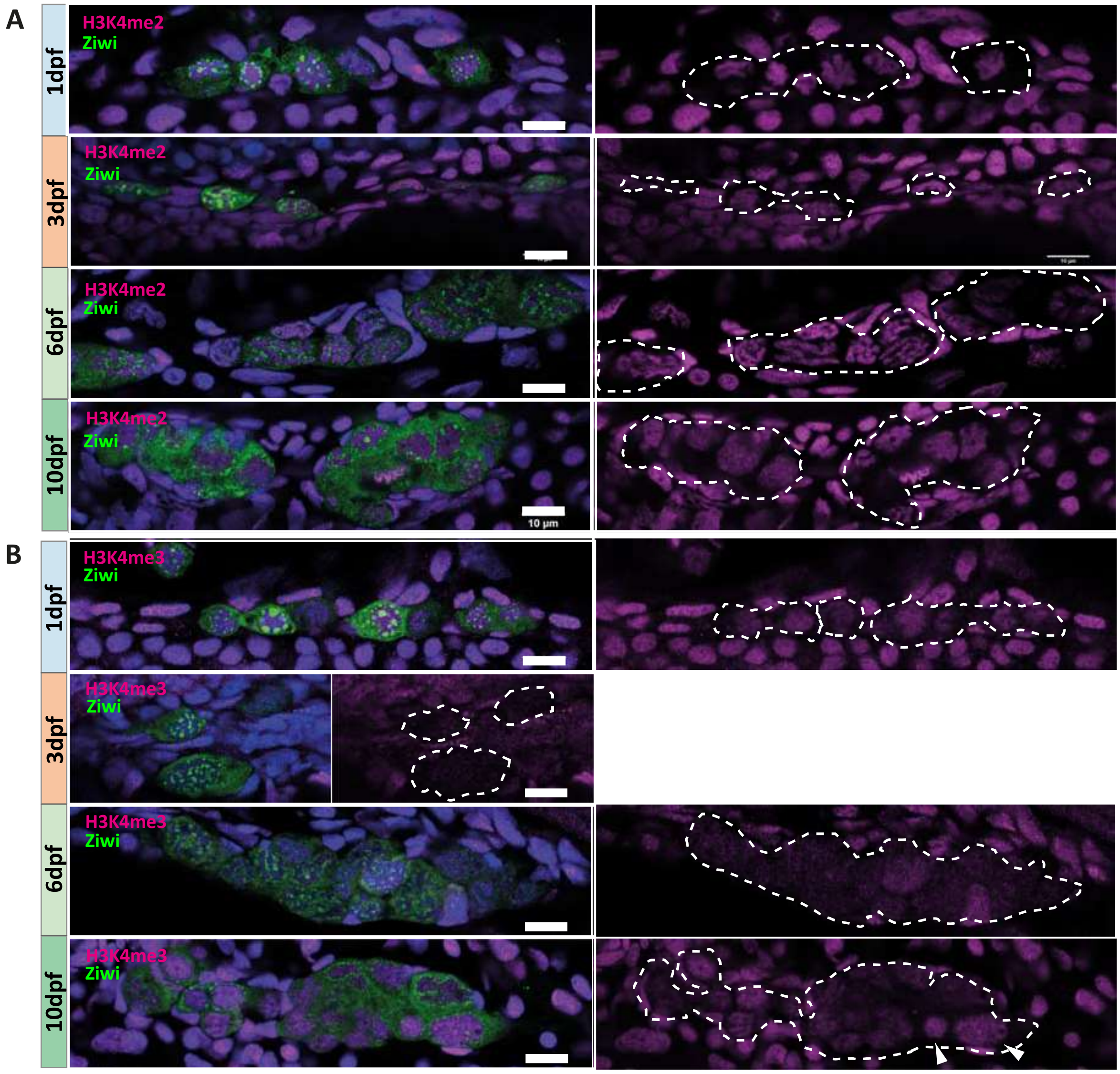

A
1dpf
H3K4me2
Ziwi
3dpf
H3K4me2
Ziwi
6dpf
H3K4me2
Ziwi
10dpf
H3K4me2
Ziwi
10 µm
B
1dpf
H3K4me3
Ziwi
3dpf
H3K4me3
Ziwi
6dpf
H3K4me3
Ziwi
10dpf
H3K4me3
Ziwi

**Figure S3**

**A**

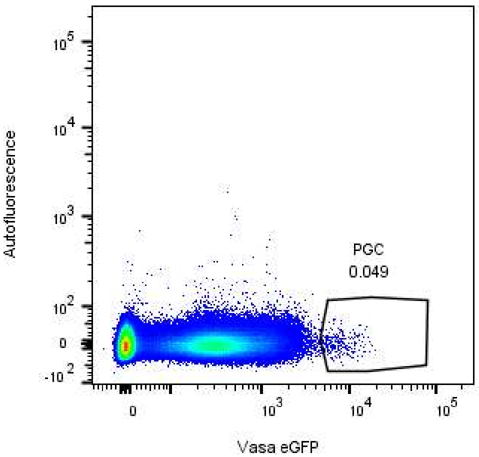


**B**

facultative tdrkh_exon

SPWVNDQLESSSRLSSSIELERADVDQNSTFKLMSSYSLI
HSTPSQNLSDLVSQILETSSSMQDPPVQSNSLTPASPLHQ
LEISSSFSSPLNVETVTSALDSFTLNDEVFLGSHYYGRNE
KTTSSASEETGSSYTLETHSENTESSNSSDGIRGVWYYLT
SSRDSSEASLSTTMPHSSTTSSSYSTEACEESDASSGSVI
ELSSDSSRSVEDAKIALRTDNQTSDLEIITLSDGSSSGSE
FVNISEEREHESETAGYLEESDPLKCDVRCQTSGFNEDVL
CNKAESGRRGSGLKNMGLKSNTGENGDESTR

**C**

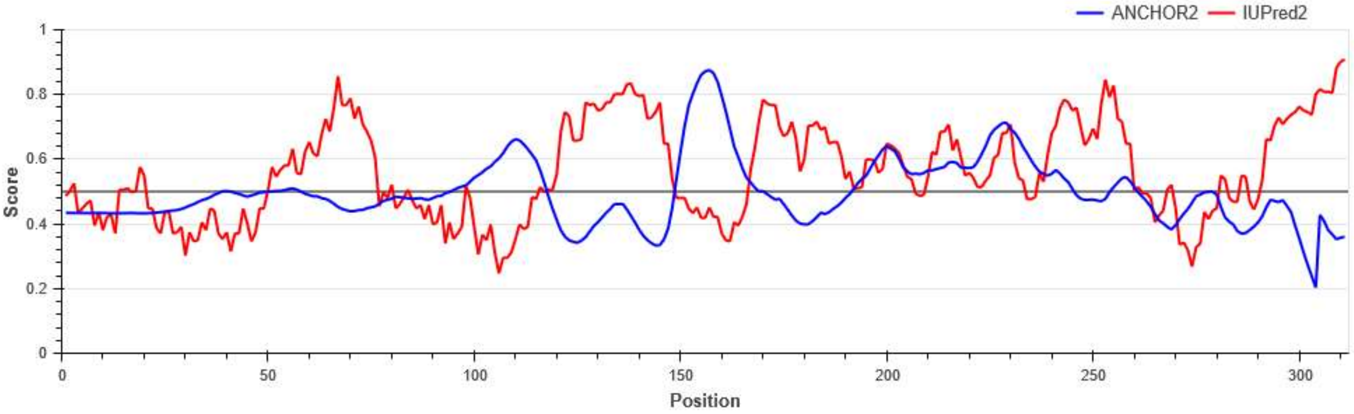


**D**

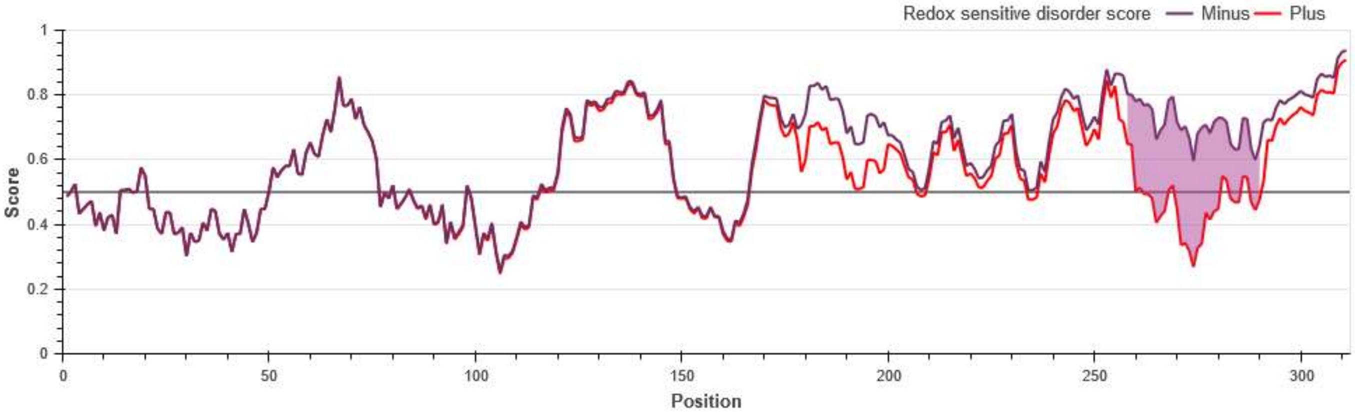

### Figure S4

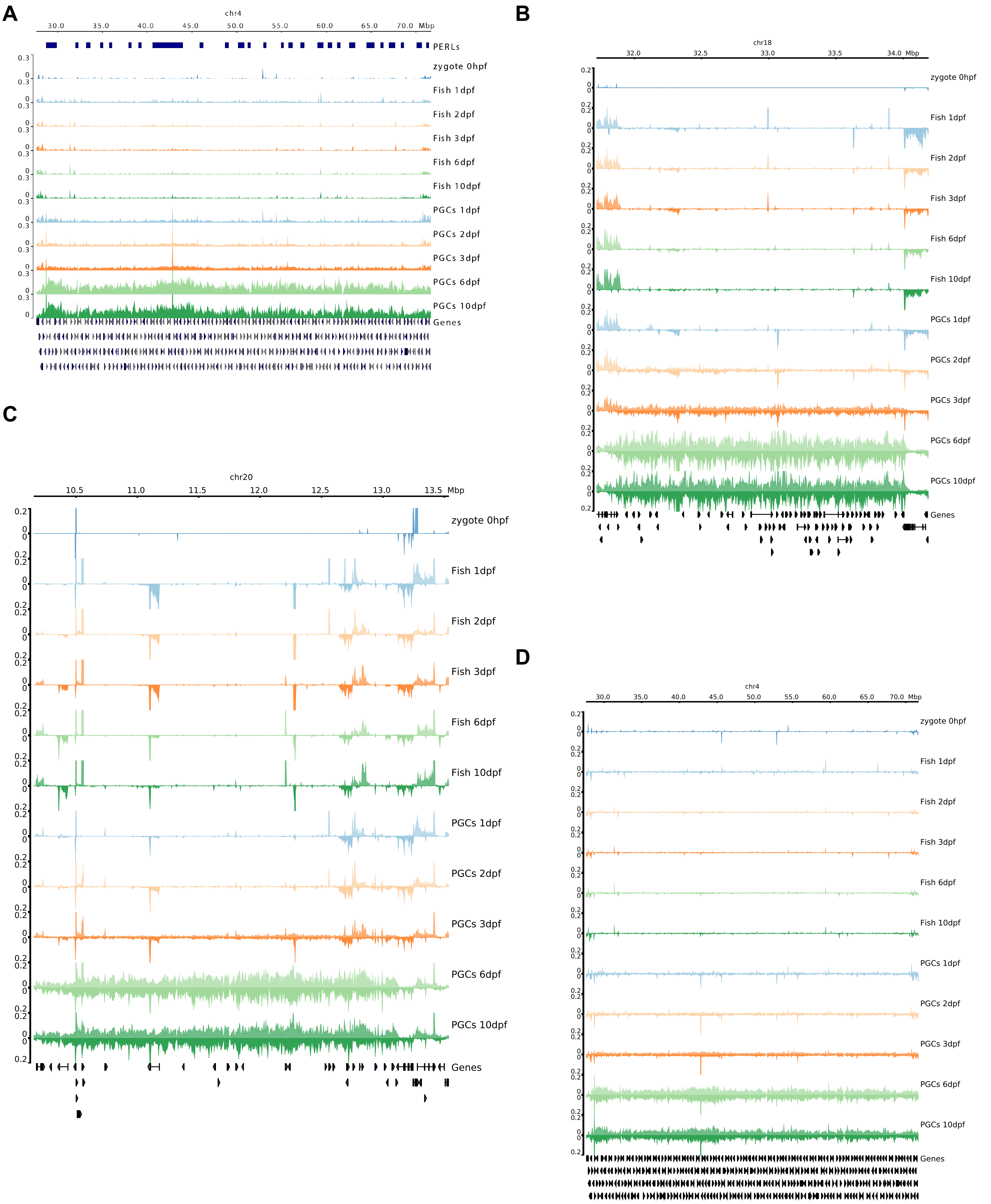

A
chr4
30.0 35.0 40.0 45.0 50.0 55.0 60.0 65.0 70.0 Mbp
PERLs
zygote 0hpf
Fish 1dpf
Fish 2dpf
Fish 3dpf
Fish 6dpf
Fish 10dpf
PGCs 1dpf
PGCs 2dpf
PGCs 3dpf
PGCs 6dpf
PGCs 10dpf
Genes
B
chr18
32.0 32.5 33.0 33.5 34.0 Mbp
zygote 0hpf
Fish 1dpf
Fish 2dpf
Fish 3dpf
Fish 6dpf
Fish 10dpf
PGCs 1dpf
PGCs 2dpf
PGCs 3dpf
PGCs 6dpf
PGCs 10dpf
Genes
C
chr20
10.5 11.0 11.5 12.0 12.5 13.0 13.5 Mbp
zygote 0hpf
Fish 1dpf
Fish 2dpf
Fish 3dpf
Fish 6dpf
Fish 10dpf
PGCs 1dpf
PGCs 2dpf
PGCs 3dpf
PGCs 6dpf
PGCs 10dpf
Genes
D
chr4
30.0 35.0 40.0 45.0 50.0 55.0 60.0 65.0 70.0 Mbp
zygote 0hpf
Fish 1dpf
Fish 2dpf
Fish 3dpf
Fish 6dpf
Fish 10dpf
PGCs 1dpf
PGCs 2dpf
PGCs 3dpf
PGCs 6dpf
PGCs 10dpf
Genes

### Figure S5

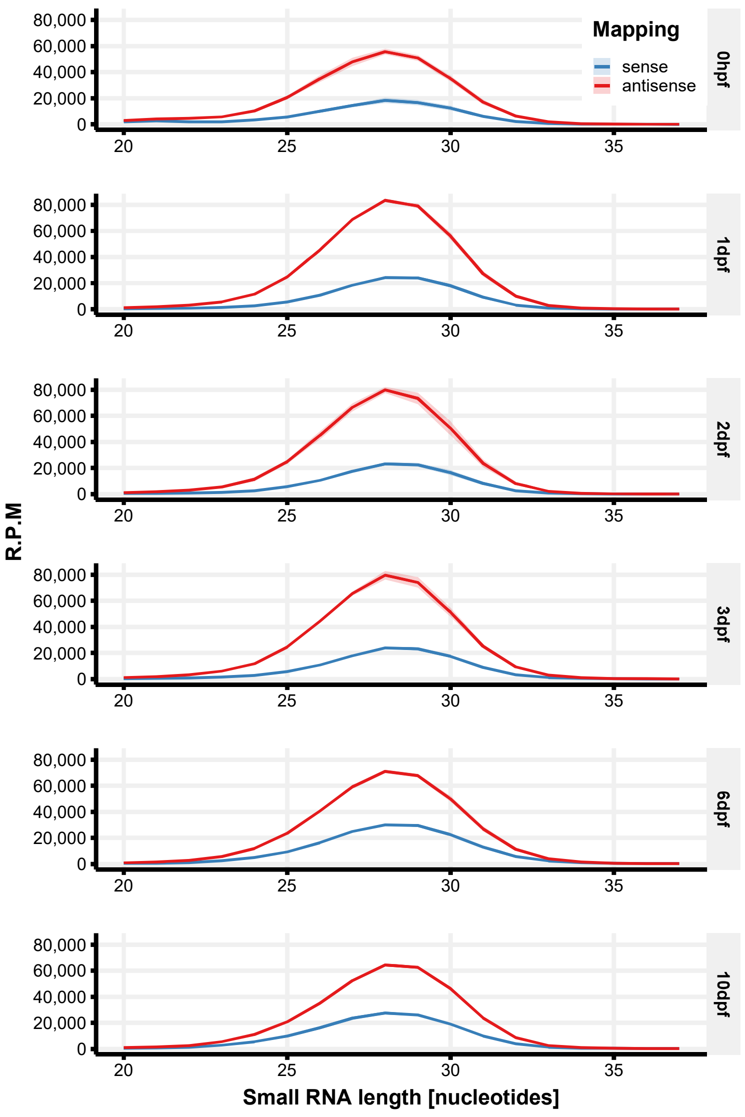

Mapping
sense
antisense
0hpf
1dpf
2dpf
3dpf
6dpf
10dpf
80,000
60,000
40,000
20,000
0
20
25
30
35
R.P.M
Small RNA length [nucleotides]
